# IFI16/AIM2 inflammasomes control Gal-9 and PVR in myeloid cells from PWH and their targeting improves immunotherapy against HIV-1

**DOI:** 10.64898/2026.01.26.701473

**Authors:** Marta Calvet-Mirabent, Ilya Tsukalov, Ildefonso Sánchez-Cerrillo, Jesús Pino-Platas, Paul Nicholas Holmes-Antón, María Agudo-Lera, Olga Popova, Raquel González, Lucio García-Fraile, José Antonio Lebrón-Romero, Manuel López-López, Sergio Serrano-Villar, Santiago Moreno, Pilar López-Cornejo, Francisco Sánchez-Madrid, Arantzazu Alfranca, Ignacio de los Santos, Enrique Martín-Gayo

## Abstract

Chronic inflammation in people with HIV-1 (PWH) is associated with low CD4/CD8 T-cell ratios and inflammasome activation, which may impair monocyte (Mo) and monocyte-derived dendritic cell (MDDC) function and limit the efficacy of myeloid-based immunotherapies. Here, we investigated the role of inflammasome sensors in regulating immune ligands for TIM3 and TIGIT immune checkpoints in Mo and MDDC from PWH and their capacity to activate HIV-1-specific CD8+ T cells. Following TLR stimulation, galectin-9 (Gal-9), PVR and IL-1β were preferentially upregulated in cells from PWH with low CD4/CD8 ratios. IFI16 and AIM2 expression was increased basally or inducible in Mo and co-localized with caspase-1 in these myeloid cells present in lymphoid tissues of HIV-1-infected humanized mice. Si-RNA or small molecule-mediated inhibition of IFI16/AIM2 reduced Gal-9 and PVR expression in MDDC and enhanced their ability to activate polyfunctional HIV-1-specific CD8+ T-cells displaying improved elimination of infected CD4+ T cells *in vitro* and *in vivo*. These findings identify IFI16 and AIM2 as key regulators of myeloid dysfunction and therapeutic targets to enhance HIV-1 immunotherapies.

## INTRODUCTION

The main barriers to achieve a cure for HIV-1 infection include the existence of persistent viral reservoirs even on antiretroviral therapy (ART), as well as the ability of the virus to efficiently evade immune responses. These factors contribute to periodic antigenic exposure in people with HIV-1 (PWH) due to punctual viral reactivation, leading to low levels of viral replication in blood and tissues ^1,2^. Such events can cause chronic inflammation and immune exhaustion, eventually resulting in tissue damage^3^, cardiovascular disorders^4^, and other Non-AIDS comorbidities^5^. In addition, HIV-1 infection has been associated with disruption of gut tissue integrity and bacterial translocation leading to the leakage of Pathogen-Associated Molecular Patterns (PAMPs) in the blood stream, and contributing to innate cell activation, systemic chronic inflammation and immune cell dysfunction in PWH on ART^6–9^. In this regard, low CD4/CD8 T cell ratio has been defined as a biomarker for a subgroup of PWH characterized by increased levels of chronic inflammation and gut dysbiosis^10–13^. Such low CD4/CD8 T cell ratios in blood from PWH can be the result of either a selective depletion of CD4+ T cells or by the expansion of CD8+ T cells caused by persistent viral particles^14,15^. Additionally, markers of T cell activation, senescence and immune exhaustion are also increased in PWH with low CD4/CD8 T cell ratios^16^. Moreover, low CD4/CD8 T cell ratios have also been associated with higher incidence of non-AIDS events in these PWH^15,17,18^. Therefore, different levels of immune exhaustion and chronic inflammation should be taken into consideration in the design of new immunotherapeutic approaches against HIV-1, including PWH with low CD4/CD8 T cell ratios.

Initial efforts to elicit HIV-1-specific T cell responses in PWH did not efficiently prevent viral rebound after treatment^19,20^, pointing to the need to fine-tune innate and adaptive antiviral immune responses. In this regard, highly functional dendritic cells (DC) prone to inducing IFN-I/TBK1 against HIV-1^21–23^ have been shown to be key promoters of effective antiviral polyfunctional HIV-1-specific CD8+ T cell responses in Elite Controllers (EC)^24–27^. In our previous studies, activation of TIGIT/TIM3 checkpoint receptors on CD8+ T cells from PWH may be a key factor determining effective response of Mo-Derived Dendritic Cells (MDDC) immunotherapies designed to mimic EC-DC ^28,29^. While a potential strategy to improve immunotherapy against HIV-1 consists in checkpoint receptor blockade, the dynamic and the molecular mechanisms regulating the expression of the ligands for these checkpoint receptors in circulating in MDDC from PWH remain unexplored.

Mo and MDDCs become activated through different innate pattern recognition receptors (PRR) which include Toll-like receptors (TLRs), innate nucleic acid sensors or inflammasome components. A potential pathogenic impact of a pre-activated state during chronic HIV-1 infection has been linked to myeloid cell dysfunction^30,31^. Several studies have reported the link between systemic immune activation and the presence of PAMPs, such as bacterial lipopolysaccharide (LPS)^8,9^, increased plasma levels of soluble CD14 (sCD14) and CD163 (sCD163), which contribute to triggering innate activation of Mo during chronic infection in PWH via TLR2, 4 ,6 ^8,32–35^. In addition, residual viral replication could also liberate viral nucleic acids that can be recognized by Mo through TLR3, TLR8^36^ or intracellular innate sensors such as RIG-I^37^. In addition to TLRs, alternative innate pathways such as the inflammasome may contribute to chronic inflammation. Previous studies have identified that HIV-1 sensing can induce the inflammasome activation^38–40^. The most classical inflammasome activation pathway involves the recognition of different pathogen or stress-associated stimuli by different intracellular sensors, such as NOD-like receptor proteins (NLRP) or nucleic acid sensors such as IFI-16 and AIM2. Activation of inflammasome sensors lead to the recruitment of the adaptor protein and the caspase-1 which form the inflammasome complex, mediating the cleavage of immature isoforms of IL-1β and IL-18, Gasdermin-D and Ninjurin-1 promoting inflammatory cell death known as pyroptosis^41–44^. HIV-1 sensing by Mo is associated with specific activation of the classical NLRP3 inflammasome and caspase-1^38,45^. In fact, inflammasome overactivation has been associated with chronic inflammation and disease progression in PWH^46,47^. However, the role of other inflammasome sensors in modulating specific functional properties in myeloid cells from PWH, and in particular expression of ligands from checkpoint receptors, has been less explored. The expression of the dsDNA inflammasome sensor AIM2 increases in HIV-1 infection, but its role in modulating the function of myeloid cells from specific PWH groups has not been studied^48,49^. Other inflammasomes such as NLRC4 and IFI16 may potentially contribute to progression of the infection by recognition and response to bacterial PAMPS^50,51^ or intracellular HIV-1 DNA^52–54^, respectively.

The present study evaluates the impact of different PAMPs in the differential expression of ligands for TIGIT (PVR) and TIM3 (galectin-9; Gal-9) checkpoint receptors in Mo and MDDC from PWH with different CD4/CD8 T cell ratios. We also identified IFI16 and AIM2 inflammasome sensors as regulators of the expression of these molecules and their pharmacological inhibition potentiates the efficacy of MDDC inducing effective HIV-1 specific CD8+ T cell responses *in vitro*. Together, our study has provided new mechanistic data that may allow to design complementary modulation approaches to improve immunotherapies against HIV-1.

## MATERIALS AND METHODS

### HIV negative and PWH Cohort blood collection and ethics statement

HIV-1 negative buffy coat samples were provided by the Centro de Transfusiones de Madrid in collaboration with the Immunology Unit from Hospital de la Princesa. Peripheral blood of a total of n=60 PWH in ART and including n=29 individuals with low (median 0.6, min 0.16-max 0.79) and n= 31 subjects with normal (median 1.36; min 1.01-max 3.73) CD4/CD8 T cell ratios provided by the Infectious Diseases Unit of Hospital Universitario de La Princesa. Clinical information for the whole PWH cohort and each patient population regarding CD4+ T cell counts, CD4+ T cell NADIR, years on ART, plasma viral load and years on ART is summarized in Supplemental Table 1. N=25 HIV negative volunteers were obtained from Centro de Transfusiones signed an informed consent granted in La Ley Orgánica 15/1999, de 13 de diciembre, de Protección de Datos de Carácter Personal and following the Helsinki declaration. All samples from PWH donors were collected at Hospital de La Princesa after receiving and signing an informed consent approved by the Ethics committee from Hospital Universitario de La Princesa (Register Number 3518) and in accordance with the Helsinki declaration. Clinical characteristics of an additional validation cohort of n=13 PWH are detailed in Supplemental Table 1 and 2.

### Peripheral blood mononuclear cell processing, Mo cell isolation and MDDC generation

Peripheral blood mononuclear cells (PBMC) were isolated by ficoll (Pancoll, PAN Biotech) gradient centrifugation from both HIV-1 negative controls and PWH. Mo were isolated from PBMC from our study cohort by the human CD14 MicroBeads kit positive immunomagnetic selection using MS Columns (Miltenyi Biotec). MDDC were generated from adhered circulating Mo and cultured for 5 days in the presence of RPMI 1640 media with 10% Fetal Bovine Serum (HyClone) supplemented with 100 IU/mL of rhGM-CSF and rhIL-4 (Prepotech).

### In vitro stimulation of Mo and MDDC with PAMPs

Isolated CD14+ Mo and MDDC from PWH on ART and from HIV-1 negative controls were cultured in the presence of 10 ng/ml of Escherichia coli lipopolysaccharides (LPS; SIGMA-ALDRICH) as a bacterial agonist for TLR4, 1.25 μg/ml of Flagellin (InvivoGen) as bacterial PAMPs activating TLR5, 2.5 μg/ml of Poly I:C (SIGMA-ALDRICH) as viral dsRNA PAMP activating TLR3/RIG-I, or 2 μg/ml of CL097 (InvivoGen) as viral ssRNA PAMP activating TLR7 and TLR8. In some experiments, MDDCs were also cultured in the presence of 1 μM Parthenolide (NF-κB and Caspase-1 inhibitor, Invivogen), 2 μM MCC950 (NLRP3 inhibitor, Invivogen), 2 μg/mL Z-VAD-FMK (Pan-caspase inhibitor, Santa Cruz Biotechnology), 6.29 μM ODN TTAGGG (Invivogen) and 1 μM BX795 (TBK-1 inhibitor, Invivogen). After incubation, culture supernatants were collected, and phenotype of Mo and MDDC was assessed as described in the Flow cytometry analysis section. Finally, in some experiments RNA was obtained from cell lysates to analyze transcriptional expression of inflammasome and proinflammatory cytokines as described in the RNA extraction and gene expression analysis section.

### RNA extraction and gene expression analysis

Total mRNA was isolated from cell lysed with RLT buffer with 1% 2-mercaptoethanol following manufacturer’s concentrations, then stored at −80°C and finally extracted using the RNeasy Micro Kit (Qiagen). cDNA was then synthesized from total RNA using QIAgen RNeasy microkit (QIAgen) and a GeneAmp PCR System 2700 (Applied Biosystems). Gene expression was analyzed by semiquantitative PCR using the SYBR Green assay GoTaq qPCR Master Mix (Promega) with 0.5 μM standardized primers (Metabion; listed on Supplemental Table 3) and 1 μl of cDNA to a total of 10 μl per reaction per duplicate. Amplification was performed either on a StepOne Plus Real-Time PCR System or QuantStudio 5 Real-Time PCR System (both from Applied Biosystems) and consisted of a first hold of 15 minutes at 95ªC, 45 cycles of 30 seconds at 95°C, 30 seconds at 60°C and 30 seconds at 72°C, and a final step of melting curve consisting of 1 minute at 95°C, 30 seconds at 55°C and 30 seconds at 95°C.

### In vitro siRNA mediated knock down of inflammasome sensors in Mo and MDDC

Aproximatelly 1.5 million of CD14+ Mo and MDDCs from PWH or HIV-1 negative controls were nucleofected with siRNA specific for either AIM2 or IFI16 intracellular inflammasome sensors (ON-TARGET plus SMARTpool Human, Dpharmacon) using the Amaxa P3 primary cell Nucleofection Kit (LONZA) in a 4D-Nucleofector Core Unit (LONZA). As a negative control, irrelevant scramble siRNA sequences were used. Nucleofected cells were cultured for 16h, then stimulated the different PAMPs for 16 additional hours, at concentrations previously mentioned. MDDC were activated with 1 μg/mL of 2’3’-c’diAM(PS)2 (Invivogen) STING agonist, 2.5 μg/mL of Poly I:C (SIGMA) or the combination of both. After incubation, phenotypical maturation of Mo and MDDC was assessed by FACS as described in the flow cytometry section.

### Functional analysis of HIV-1 specific T cell responses in vitro

MDDCs were activated with 1 μg/mL of 2’3’-c’diAM(PS)2 (Invivogen) STING agonist and 2.5 μg/mL of Poly I:C (SIGMA) in the presence of DMSO or 5μg/ml Gag-peptides alone for 16h. Subsequently, activated MDDC were treated for 3h with either DMSO or 2 μg/mL Z-VAD-FMK (Santa Cruz Biotechnology), 6.29 μM ODN TTAGGG (Invivogen) soluble or loaded into liposome-based nanosystems (LBN) (see Liposome-based nanosystems preparation section). Autologous CD8+ T cells were isolated according to the manufacturer’s protocol (MojoSort) and cocultured with treated MDDCs in presence of IL-2 (50IU/ml) for 2 hours and then 16 hours in presence of 2.5 μg/ml Brefeldin A (MERCK), 2.5 μg/ml Monensin (Sigma) and anti-CD107a (1:600 dilution). Expression of CD107a, IFN-γ and TNFα was analyzed by FACS (see flow cytometry section). Membrane molecules were stained at 1:50 dilution and intracellular at 1:100 dilution. As a positive control of degranulation and cytokine secretion 0.025 μg/ml of PMA (MERCK) and 0.125 μg/ml of Ionomycin (Peprotech) were used.

### Killing assays

As previously stated, the MDDCs activated with of 2’3’-c’diAM(PS)2, Poly I:C and Gag-peptide for 16h and subsequently treated with Z-VAD-FMK, ODN TTAGGG or DMSO for 3 hours were cocultured with autologous CD4+ and CD8+ T lymphocytes isolated according to the manufacturer’s protocol (Invitrogen, MojoSort) in presence of IL-2, Raltergravir (Selleck Chemical) and Romidepsin (Selleck Chemical) for 16 hours. Expression of intracellular p24 in cultured CD4+ T cells was determined by FACS (see flow cytometry section).

### Flow cytometry analysis

Cells were harvested and block with 20 μg/ml of human gamma-globulins (SIGMA-ALDRICH) was performed to avoid unspecific antibody binding to Fc Receptors. Ghost Dye Red 780 (Tombo biosciences) or LIVE/DEAD Fixable Yellow 405 (Invitrogen) viability dyes were included to determine cell viability, depending on the panel configuration. In the case of panels including intracellular staining, after labelling membrane molecules, cells were washed and incubated for 30 minutes at 4°C with Fixation buffer 1x (Biolegend) and then incubated with intracellular antibodies included in 100 μl of Permeabilization buffer (Biolegend).

Human PBMC were analyzed on a FACS Canto II or on a BD FACS Lyric cytometer (BD Biosciences) at the Immunology Unit of Hospital Universitario de La Princesa (Madrid, Spain). Briefly, for analysis of myeloid cell activation after PAMP incubation mAbs against PD-L1, PVR, Gal-9, HLA-DR, CD40 and CD86 were used as membrane markers and TNFα, IL-1β and IL-6 as intracellular markers. For analysis of *in vitro* CD8+ T cell activation, we used Ghost Red viability Dye 780 (Tombo biosciences), anti-CD3-Pacific Blue (Immunostep), anti-CD8-PeCy7 (BD), anti-CD107a-APC (Biolegend); anti-IFN-γ-FITC (BD) and anti-TNFα-PerCP (Biolegend). To analyze *in vitro* functional capacity of CD8+ T cells to eliminate HIV-1 infected cells, staining with Ghost Dye Red 780 (Tombo biosciences), anti-CD3 FITC (Biolegend), anti-P24 PE (Beckman Coulter), anti-CD8 PeCy7 and anti-CD4 Pacific Blue (Biolegend) mAbs were used. Further information about antibody clones and commercial sources is detailed in Supplemental Table 4.

FACS analysis of in vivo experiments was performed using a panel that included Ghost Red viability Dye 780 (Tombo biosciences), anti-CD3-BV605 (Biolegend), anti-CD8-BV786 (Biolegend), anti-CD45-V500 (Biolegend) and anti-p24-PE (Beckman Coulter) (Supplemental Table 4).

### Microscopy immunofluorescence analysis of inflammasome activation in Mo and MDDC

The colocalization between IFI16, AIM2 and Caspase-1 inflammasome components was carried out by immunofluorescence from isolated Mo or MDDC cultured for 16 h in presence of Poly I:C or media and stained with either a rabbit anti-human IFI16 (abcam) or a rabbit anti-human AIM2 (abcam) in combination with goat anti-human Caspase-1 (Bio-Techne RyD Systems). Secondary antibodies included donkey anti-goat AlexaFluor™ 568 (Invitrogen) and donkey anti-rabbit AlexaFluor™ 488 (Invitrogen) secondary antibodies, supplemented with DAPI (Thermofisher). Images were acquired in a Leica TCS SP5 (Leica MS GmbH) confocal microscope and co-localization was analyzed using ImageJ software (version 1.54r, including LabKit plugin) (Supplemental Table 5).

### Histological analysis of BLT spleen tissue

Spleens from humanized BLT mouse infected with JRCSF HIV-1 and displaying high viremia were embedded in paraffin and segmented in 5 μm fragments for histological analysis. Tissue section deparaffinization, hydration and antigen retrieval were performed prior to antibody staining. Tissue slices were stained with primary antibodies rabbit anti-human IFI16 (abcam), rabbit anti-human PVR (Cell signaling), goat anti-human Caspase-1 (Bio-Techne RyD Systems) and mouse anti-human CD14 (abcam) using a basic antigen retrieval buffer (Dako). Secondary antibodies included donkey anti-rabbit AlexaFluor 488 (Invitrogen), donkey anti-mouse AlexaFluor 647 (Invitrogen) and donkey anti-goat AlexaFluor 568 (Invitrogen), supplemented with DAPI. Images were obtained with THUNDER microscope (Leica MS) Las X software (version 10.1) and analyzed with QuPath software^55^ (version 0.6.0) (Supplemental Table 5).

### Quantitative analyses of soluble biomarkers

Concentration of gram-negative bacterial endotoxin in plasma was quantified from plasma in triplicate using the Limulus Amebocyte Lysate Kinetic-QCL assay (LONZA). Kinetics of endotoxin were measured at 405 nm every 150 seconds for 1 hour and 40 minutes in a GloMax Discover instrument (Promega). Reaction time for each well was considered as the time where the increase in absorbance from initial measured absorbance was 0.2.

Plasma levels of soluble CD14, IL-6 and IL-1β were determined using the Human CD14 DuoSet ELISA, the Human IL-6 DuoSet ELISA kits (both from R&D systems) or the Quantikine ELISA Human IL- 1β/IL-1F2 kit (R&D systems), respectively. Optical density was measured at 450 nm with correction at 560 nm in a GloMax Discover instrument (Promega).

### Liposome-based nanosystems preparation

Liposome-based nanosystems (LBN) were prepared from 1 mg/ml of L-α-phosphatidylcholine from egg yolk (PC) (Sigma Aldrich), cholesterol (CHO) (Sigma Aldrich) and anti-CD64 mAb (Biolegend) using the thin-film hydration method.^56^ A mass ratio of PC:CHO:anti-CD64mAb of 1.0:0.5:0.0025, respectively, was dissolved in 1 mL of a mixture chloroform:ethanol (2:1 v/v) and sonicated for 2 min (JP Selecta Ultrasounds, 200 W and 50 kHz) at room temperature. The organic solvent was evaporated in a centrifuge concentrator (Eppendorf Concentrator plus, Hamburg, Germany) for 60 min at room temperature, and the resulting lipid bilayer was stored at −81°C for 24 h to avoid phospholipid degradation.

Consequently, lipid bilayers were hydrated with 2 mL of PBS buffer at pH=7.2 and introduced to a process of 10 cycles of vigorous agitation, consisting of a vortex process for 3 min at 1200 rpm followed by a 2 min ultrasound process. Hydration then continues with a vigorous vortex agitation process for 2 h at 1200 rpm in darkness. All the nano-formulations prepared were extruded with a miniextruder Avanti Polar Lipids and polycarbonate membranes (100 and 200 nm diameters) from Whatman were used in this process. The extrusion was repeated 10 times. Morphology of resulting anti-CD64mAb decorated LBN was tested by transmission electron microscopy, by depositing part of the LNP preparation on a copper grid coated with a carbon film. Samples were dried in the grid at room temperature and in the dark. The images were viewed using a Talos S200 high-resolution transmission electron microscope.

Binding of LBN to Mo was assessed *in vitro*, using LBN loaded with 0.9 mM of 5(6)-carboxyfluorescein (CFCL) of either naked or 2.5 or 10 μg of anti-CD64 mAb decorated LBN. Specific binding of 15 μg/ml of either naked or 2.5 or 10 μg anti-CD64 mAb decorated LBN to Mo was evaluated by FACS. For the preparation of LBN containing Z-VAD-FMK, the caspase inhibitor was added to the lipid bilayer at a different mass ratio 1.33:0.67:0.0033 (PC:CHO:anti-CD64) ratios, corresponding to a final 0.2 mg/mL Z-VAD-FMK concentration.

### In vivo assays

Immunodeficient NOD/SCID IL2R-y−/− (NSG) were acquired from Charles River and used for the *in vivo* experiments using the humanized mouse outgrowth assay (mVOA) model of HIV-1 infection. Animals were housed at the BSL3 facility from Centro de Biología Molecular Severo Ochoa (CBMSO) in accordance with the institutiońs animal care standard. Experimental procedures were reviewed and approved by the local ethics committee and were in agreement with the EU Directive 86/609/EEC, Recommendation 2007/526/ec and real Decreto 53/2013.

Prior to the in vivo experiments, adhered circulating Mo from PWH were cultured for 5 days in the presence of RPMI 1640 media with 10% Fetal Bovine Serum (HyClone) supplemented with 100 IU/mL of rhGM-CSF and rhIL-4 (Prepotech) to generate MDDCs. 24h prior injection of cells from PWH, a total of *n* = 22 NSG mice (2 independent experiments) were irradiated with 1.5 Gr to facilitate human cell engraftment. In parallel, MDDCs were activated with 1 μg/mL of 2’3’-c’diAM(PS)2 (Invivogen) STING agonist, 2 μg/mL of Poly I:C (SIGMA) and loaded with 5μg/ml Gag-peptides for 16h. Also, 24h prior to infusion CD4 and CD8 T cells isolated by negative magnetic sorting from PBMC from the same PWH donor according to the manufacturer’s protocol (Invitrogen, MojoSort) and activated with ImmunoCult™ Human CD3/CD28 T Cell Activator (Stemcell Technologies) and in the presence of 50IU/mL IL-2 for 16h. Subsequently, mice were intravenously injected with a range of 200,000-1,000,000 of CD4+ T cells alone (n=7) or in combination with 100,000-300,000 MDDCs and 200,000-400,000 CD8+ T cells per animal. Two independent experiments were performed using two different PWH donors. In addition, NSG mice were injected intraperitoneally in 200 μl of either PBS (n=7) or empty (n=8) or Z-VAD-FMK loaded (n=7) CD64-decorated LBN at day 0 and repeated after 3 days. After 7 days from the first treatment animals were sacrificed and spleen was extracted, to perform FACS analysis of proportions of HIV-1 p24+ cells included in CD45+ human CD4+ T cells.

### Statistical analysis

Statistical significance of differences between paired conditions were assessed either using non-parametric two-tail Wilcoxon matched-pairs signed-rank test or multiple comparison Friedman test followed by post-hoc Dunn’s correction method. Non- parametric two-tail Mann-Whitney U test or Kruskal-Wallis multiple comparison test followed by post-hoc Dunn’s correction method to assess significant differences between independent samples. Individual correlations and correlation networks were both generated from individual non-parametric Spearman correlations. All these statistical tests were performed using GraphPad Prism software version 8.

## RESULTS

### Induction of TIGIT and TIM3 ligands in Mo from donors with low CD4/CD8 T cell ratios

A cohort of n=60 PWH on ART for at least 1 year was recruited to the study, which was stratified in two subgroups of individuals with different low (<0.8 n=29) or normal (>1; n=31) CD4/CD8 T cell ratios in blood (Supplemental Table 1). Furthermore, while the majority of PWH included in our cohort was characterized by undetectable levels of plasma viral load (median 20 copies/μl VL >0.8 median 20 copies/ul VL<0.8 ratios), PWH with low ratios significantly contained higher proportions of individuals with residual pVL below 200 copies/mL (Supplemental Table 1). However, no significant association between pVL and other clinical parameters was observed within the PWH with low CD4/CD8 T cell ratio group (Supplemental Figure 1A-B). As expected, when considering all PWH, we observed a positive correlation between CD4+ T cell counts at sample collection or at diagnosis (NADIR) as well as years on ART treatment with higher CD4/CD8 ratios (Supplemental Figure 1A, left). We also observed a significant correlation between years on ART and age, which was significant in the low CD4/CD8 ratio PWH group (Supplemental Figure 1A). Therefore, we asked whether parameters of systemic inflammation and the years under ART could be associated with the two CD4/CD8 T cell ratio PWH groups. To address this, we first evaluated whether plasma levels of different proinflammatory cytokines (IL-6, IL-1β) and soluble markers associated to bacterial translocation and microbiota disruption (soluble CD14 and endotoxin) were associated with clinical parameters at collection or at infection in a small group of PWH including n= 12 normal and n=8 low ratio individuals from our cohort. IL-1β plasma levels tended to be higher (p=0.0568) in PWH with low CD4/CD8 T cell ratio compared to subjects with normal CD4/CD8 T cell values (Figure 1A). In contrast, IL- 6 was only detected in the plasma of three PWH with low CD4/CD8 T cell ratio and no significant differences in plasma sCD14, endotoxin and tryptophan levels were found in PWH groups from our study cohorts (Supplemental Figure 1B). Although no direct association with bacterial translocation was observed in the two groups of PWH, IL-1β plasma levels also positively correlated with plasma levels of sCD14 (Figure 1B). Overall, these data suggest that higher IL-1β plasma levels support a higher inflammatory state in PWH with low CD4/CD8 T cell ratio and less years under ART.

**Figure 1.**
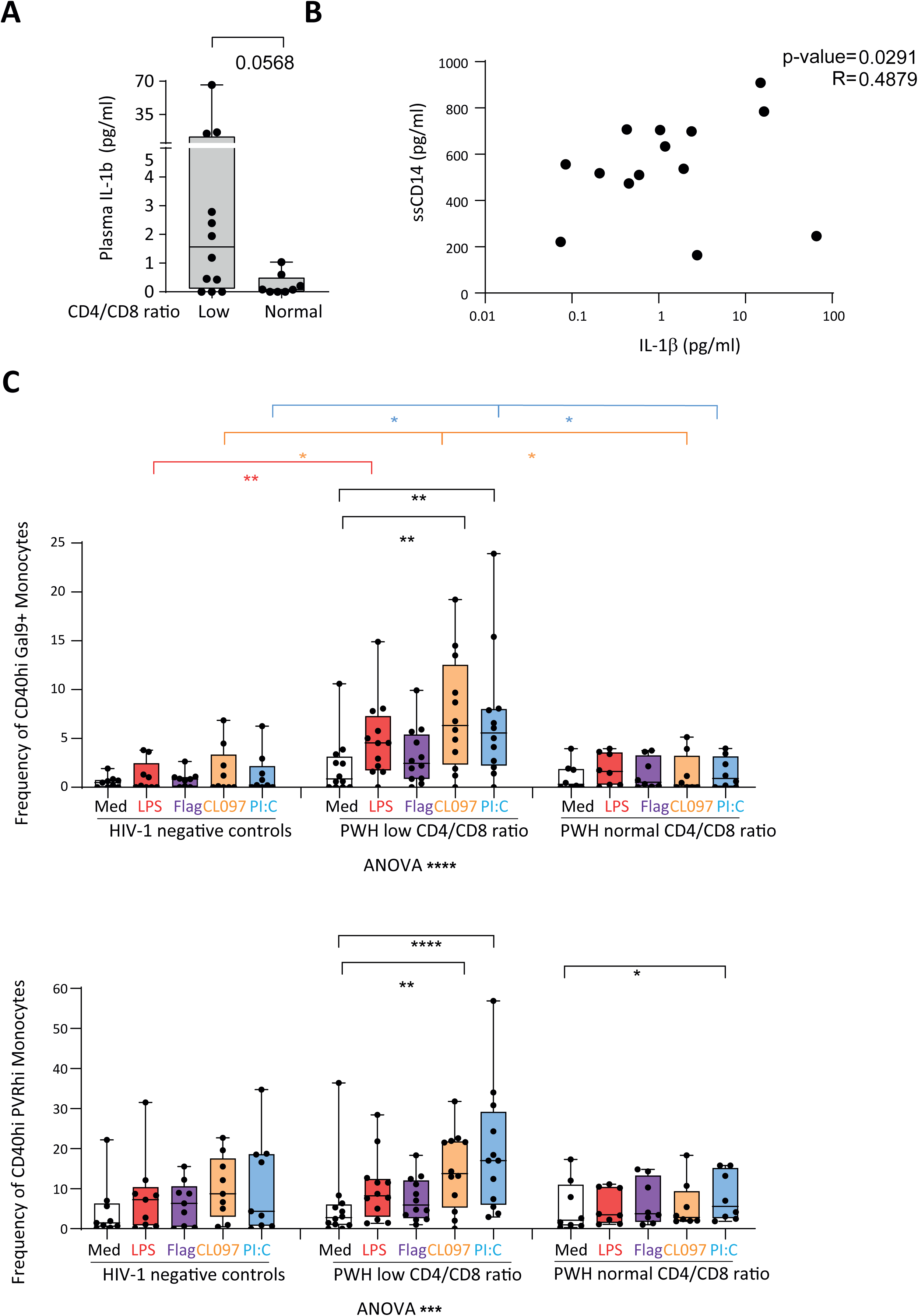
Plasma markers of inflammation and bacterial translocation in PWH. (A): ELISA analysis of plasma concentration of IL-1β, comparing PWH with normal CD4/CD8 T cell ratio (defined as ≥1) and low CD4/CD8 T cell ratio (defined as < 0.8). Statistical significance was calculated using two-tailed Wilcoxon test. (B): Spearman correlation between plasma concentrations of soluble CD14 (ssCD14) and IL-1β. P and R values are shown (C): Expression of Gal-9 (Gal9; higher plot) or PVRhi (lower plot) in activated CD40hi in Mo from HIV- 1-negative controls, and PWH with low CD4/CD8 T cell ratio or normal CD4/CD8 T cell ratio after 16-hour stimulation with PAMPs mimicking bacterial translocation (LPS, red; Flagellin, Flag, purple) or viral replication (CL097, ssRNA, orange; Poly I:C, dsRNA, blue). Statistical significance was calculated using Kruskal-Wallis test for comparison of stimulation with the same PAMP between groups (*p<0.05; **p<0.01), and One-Way ANOVA Friedman test with Dunn’s multiple comparison test for comparison within the same group (*p<0.05; **p<0.01; ***p<0.001; ****p<0.0001).

We next evaluated in these donors and HIV-1 negative donors (HD), the surface co-expression of ligands for PD-1 (PD-L1), TIGIT (PVR/CD155) and TIM-3 (Gal-9), together with the maturation marker CD40, in Mo either at baseline or exposed to different PAMPs that mimic bacterial translocation (LPS and flagellin) and viral replication (ssRNA analog CL097 and dsRNA analog Poly :IC) (Supplemental Figure 1C). Unexpectedly, we found that Mo from PWH with lower CD4/CD8 T cell ratios tended to display lower basal levels of CD40 than HD or PWH with normal CD4/CD8 T cell ratios, but they were able to significantly increase the expression of this molecule upon stimulation with most tested PAMPs (Supplemental Figure 1D).

PVR and Gal-9 expression was significantly increased on CD40hi Mo after treatment with CL097, Poly I:C and LPS, and to a lower extent in the presence of Flagellin, both in PWH and HD control individuals (Figure 1C, Supplemental Figure 1C). However, the proportions of CD40hi Mo expressing Gal-9 and PVR were significantly higher in PWH with low CD4/CD8 T cell ratios compared to HD and/or PWH with normal CD4/CD8 T cell ratios after exposure to Poly I:C or CL097 viral PAMPs (Figure 1C). Notably, to rule out a potential effect of residual low pVL in PWH with low CD4/CD8 ratios we confirmed that higher upregulation of PVR and Gal-9 was also observed in Mo from aviremic fully suppressed PWH with low CD4/CD8 T cell ratios (Supplemental Figure 1E-F). Of note, proportions of Mo expressing PVR and Gal-9 in Mo from PWH with normal CD4/CD8 ratio was similar to cells from HIV negative control individuals after PAMP stimulation (Figure 1C). Similar results were observed for PD-L1, although differences in the expression of this ligand were less obvious between PWH with different CD4/CD8 T cell ratios (Supplemental Figure 1G). Overall, our data indicate a higher inducibility of expression of ligands of checkpoint inhibitory receptors TIGIT and TIM3 in Mo from PWH with low CD4/CD8 T cell ratios after exposure to viral and bacterial PAMP stimulation.

### Increased expression of IFI16/AIM2 inflammasomes in Mo from PWH with low CD4/CD8 T cell ratios

To better dissect potential mechanisms that drive the enhanced upregulation of PVR and Gal-9 in Mo from PWH with low CD4/CD8 T cell ratios, we first evaluated transcription of pro-inflammatory cytokines (IL-1β, IL-6 and TNFα) and IFN-β in response to PAMPs in these cells, compared to Mo from PWH with normal ratios and HD. As expected, LPS, CL097 and Poly I:C significantly induced transcription of TNFα and IL-6 (Supplemental Figure 2A). Induction of TNFα transcription upon TLR stimulation was significantly higher in Mo from all PWH groups and HIV-1 negative donors, but it was more significant in cells from PWH with low and normal CD4/CD8 ratio groups (Supplemental Figure 2A). Importantly, IFNβ transcription was upregulated by LPS in the HIV-1 negative control and PWH groups, but its induction in response to CL097 and Poly I:C tended to be preferentially observed in Mo from PWH with low CD4/CD8 T cell ratios, although it was not significant (Supplemental Figure 2A). However, induction of IL-1β and IL-6 transcripts were significantly higher in response to LPS (p-value 0.0002 and 0.0001 respectively) and Poly I:C (p-value 0.0029 and 0.0029 respectively) specifically in Mo from PWH with low CD4/CD8 T cell ratios compared to baseline levels, in contrast to PWH donors with normal ratios exhibited lower level of induction for most tested PAMPs (Figure 2A, Supplemental Figure 2A). Therefore, these data suggest that Mo from PWH with low CD4/CD8 ratios are characterized by a higher inflammatory profile. Next, we asked whether the secreted isoform of IL-1β could be differentially detected in supernatants of Mo from the different PWH groups after the exposure with different PAMPs. From all PAMPs evaluated, CL097 most consistently induced significant increase in IL-1β concentrations in culture supernatants from cells in all PWH and the HIV-1 negative control groups (Figure 2B). In contrast, LPS and Poly I:C stimulation differentially induced a significant increase of secreted IL-1β in culture supernatants of Mo from PWH, and more significantly in the case of cells from individuals with low CD4/CD8 ratios (Figure 2B).

**Figure 2.**
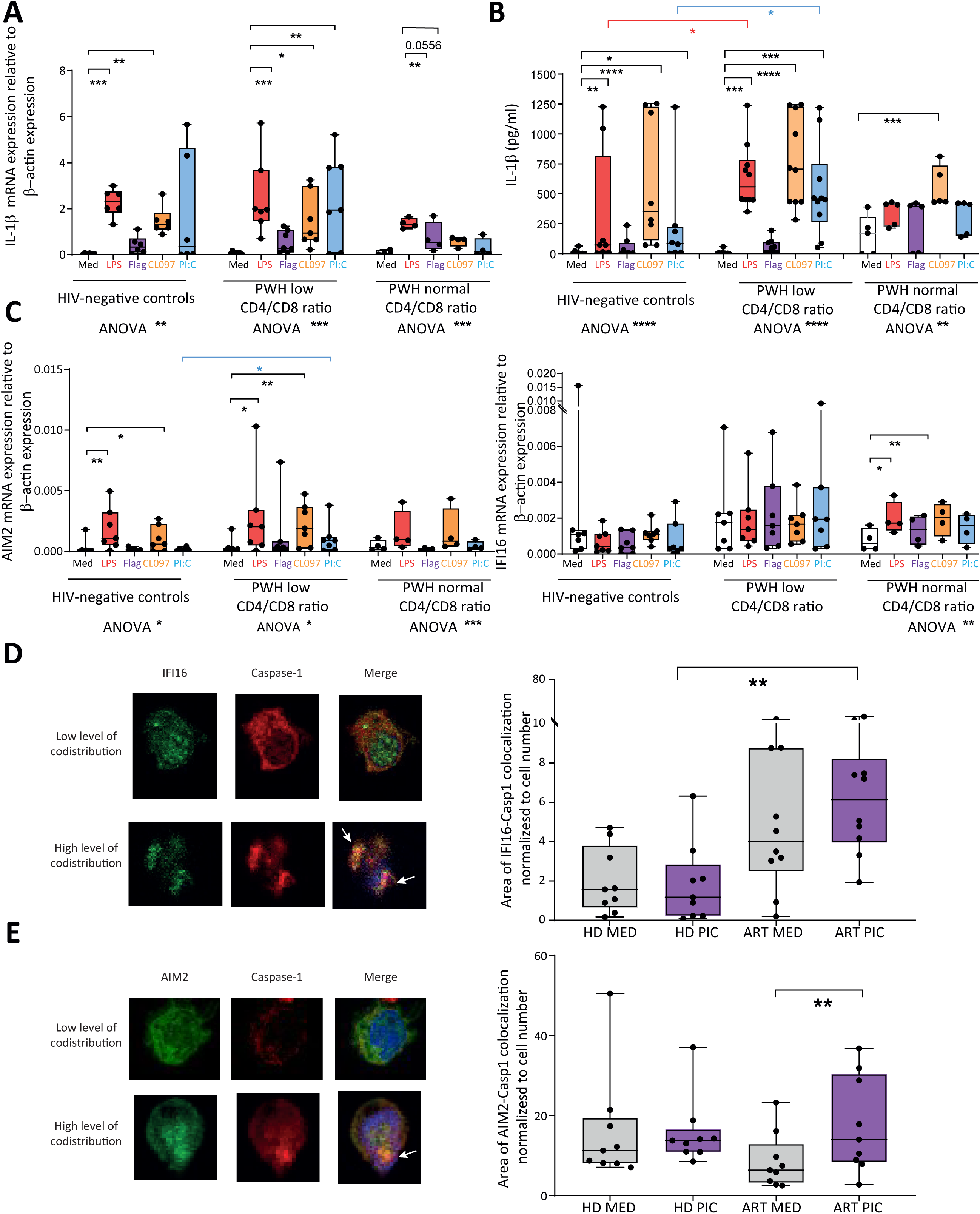
Analysis of inflammasome activity in Mo from PWH with low and normal CD4/CD8 T cell ratios. (A,C): RT-qPCR analysis of IL1β (A), AIM2 (C, left) and IFI16 (C, right) transcriptional expression normalized to β-actin mRNA levels in Mo from HIV-1 negative controls, and PWH with low CD4/CD8 T cell ratio or normal CD4/CD8 T cell ratio after 16-hour stimulation with PAMPs mimicking bacterial translocation (LPS, red; Flagellin, Flag, purple) or viral replication (CL097, ssRNA, orange; Poly I:C, dsRNA, blue). (B): ELISA analysis of IL-1β concentration (pg/mL) in culture supernatants of Mo from HIV negative and PWH with low and normal CD4/CD8 T cell ratios cultured in the conditions previously described for (A,C). (D,E): Confocal microscopy immunofluorescence analysis of co-expression of caspase-1 and IFI16 (D) or AIM2 (E) in Mo from HIV negative (HD) and PWH (ART) donors. Representative zoomed 63X magnification images showing individual or co-distribution of these inflammasome proteins are shown on the left. Statistical significance was calculated using Kruskal-Wallis for comparison of stimulation with the same PAMP between groups (*p<0.05; **p<0.01; ***p<0.001; ****p<0.0001), and One-Way ANOVA Friedman test with Dunn’s multiple comparison test for comparison within the same group (*p<0.05; **p<0.01; ***p<0.001; ****p<0.0001). Statistical significance for the colocalization assays was calculated using two-tailed Mann-Whitney in HD-ART comparison and Wilcoxon test for same group comparisons (**p<0.01).

Since secretion of active IL-1β is the functional consequence of the activation of the inflammasome complex^57^, we evaluated the transcriptional expression of different inflammasome sensors in Mo from the different PWH subgroups and HIV-1 negative controls by RT-qPCR. Among all potential inflammasome sensors, we selected NLRP3 as the most studied canonical inflammasome that can be activated by TLR signaling that has been involved in the loss of CD4+ T cell by pyroptosis during HIV-1 infection^58^. We also analyzed expression of NLRC4, which can also be induced in response to bacterial PAMPs^59,60^. However, while a tendency to lower NLRC4 and higher NLRP3 expression was observed in Mo from PWH, no significant differences were observed in baseline transcription of these sensors in Mo from HIV-1 negative controls and the different PWH subgroups (Supplemental Figure 2B). We then analyzed expression of alternative inflammasome sensors associated with sensing of intracellular nucleic acids, such as AIM2 and IFI16. AIM2 transcriptional levels increased in response to LPS and CL097 in cells from all PWH and HD groups but were most significantly induced in Mo from PWH with low CD4/CD8 ratios (Figure 2C, left). Importantly, IFI16 transcriptional levels tended to be basally higher in Mo from low CD4/CD8 T cell ratios and remained stable after PAMP stimulation (Figure 2C, right). In contrast, basal levels of IFI16 in PWH with normal CD4/CD8 T cell ratios were similar to those present in HIV-1 negative controls, but LPS and CL097 stimulation significantly induced upregulation of the IFI16 transcription specifically in these PWH individuals, (Figure 2C right). Therefore, IFI16 and AIM2 sensors may differentially promote inflammasome activation in Mo from PWH with low CD4/CD8 ratios in response to PAMP stimulation. To evaluate this possibility, we performed an analysis of expression and co-distribution of IFI-16 or AIM2 and caspase-1 in Mo from HD and CD4/CD8 low ratio PWH at baseline and after Poly I:C stimulation. IFI-16 mainly distributed in the nucleus at baseline condition but adopted a cytoplasmic location upon activation (Figure 2D). Notably, a higher level of co-localization of cytoplasmic IFI-16 and caspase-1 was observed in Mo from PWH with low CD4/CD8 T cell ratios compared to cells from HD in response to Poly I:C confirming the inflammasome assembling (Figure 2D). In contrast, AIM2 showed a cytoplasmic distribution at both basal and stimulation conditions, and no obvious differences in basal caspase-1-AIM2 co-localization was observed between Mo from PWH and HD at baseline. However, upon Poly I:C stimulation a significant increase of AIM2-caspase 1 colocalization was specifically observed in Mo from low CD4/CD8 ratio PWH (Figure 2E). Therefore, the data indicate higher level of IFI-16 and AIM2 inflammasome activation in response to Poly I:C in Mo from PWH with low CD4/CD8 ratios.

### Higher expression of IFI-16/AIM2 inflammasomes and caspase in Mo from lymphoid tissue of HIV-1 infected humanized mice

To determine the physiological relevance and the potential connection between activation of the IFI16/AIM2 inflammasomes and expression of TIGIT and TIM3 ligands, we took advantage of available tissue from humanized BLT mice infected with HIV-1 with different levels of plasma viral loads. As shown in Figure 3A (upper plot), IFI-16 expression was more concentrated in the white pulp areas where HIV-1 infected p24+ cells were present, and also basally in these areas in uninfected mice (Supplemental Figure 3A-C). However, IFI16+ cells could also be found in red pulp of infected animals (Figure 3A, upper plot). High co-expression of IFI-16 and Caspase-1 was also present in the white pulp of spleen from HIV-1 infected humanized BLT mice (Figure 3A, middle plot) and such colocalization was more obvious in tissue from animals with higher viral loads as compared to tissue from uninfected animals (Supplemental Figure 3A, B). Interestingly, IFI16+ Caspase 1+ CD14+ Monocytes were also preferentially located in the white pulp from infected animals (Figure 3A, bottom plot). Interestingly, although detection of AIM2 showed opposite patterns than IFI16 with higher number of positive cells in the spleen red pulp (Figure 3B, upper plot), Caspase1+ AIM2+ CD14+ Mo also tended to preferentially locate in the white pulp of the spleen from HIV-1 infected hBLT mice (Figure 3B lower plot, Supplemental Figure 3B). Finally, co-expression of caspase-1 and Gal-9 was also observed in Mo mainly localized in the white pulp from the lymphoid tissue of viremic HIV-1 infected humanized mice (Figure 3C). Less obvious, co-expression of PVR and caspase-1 was observed in Mo present in these tissues (Supplemental Figure 3E). Therefore, our data suggest that high level of IFI16/AIM2 inflammasome activation in lymphoid tissue during progressive HIV-1 infection, and these patterns associate with increased expression of PVR in these tissues.

**Figure 3.**
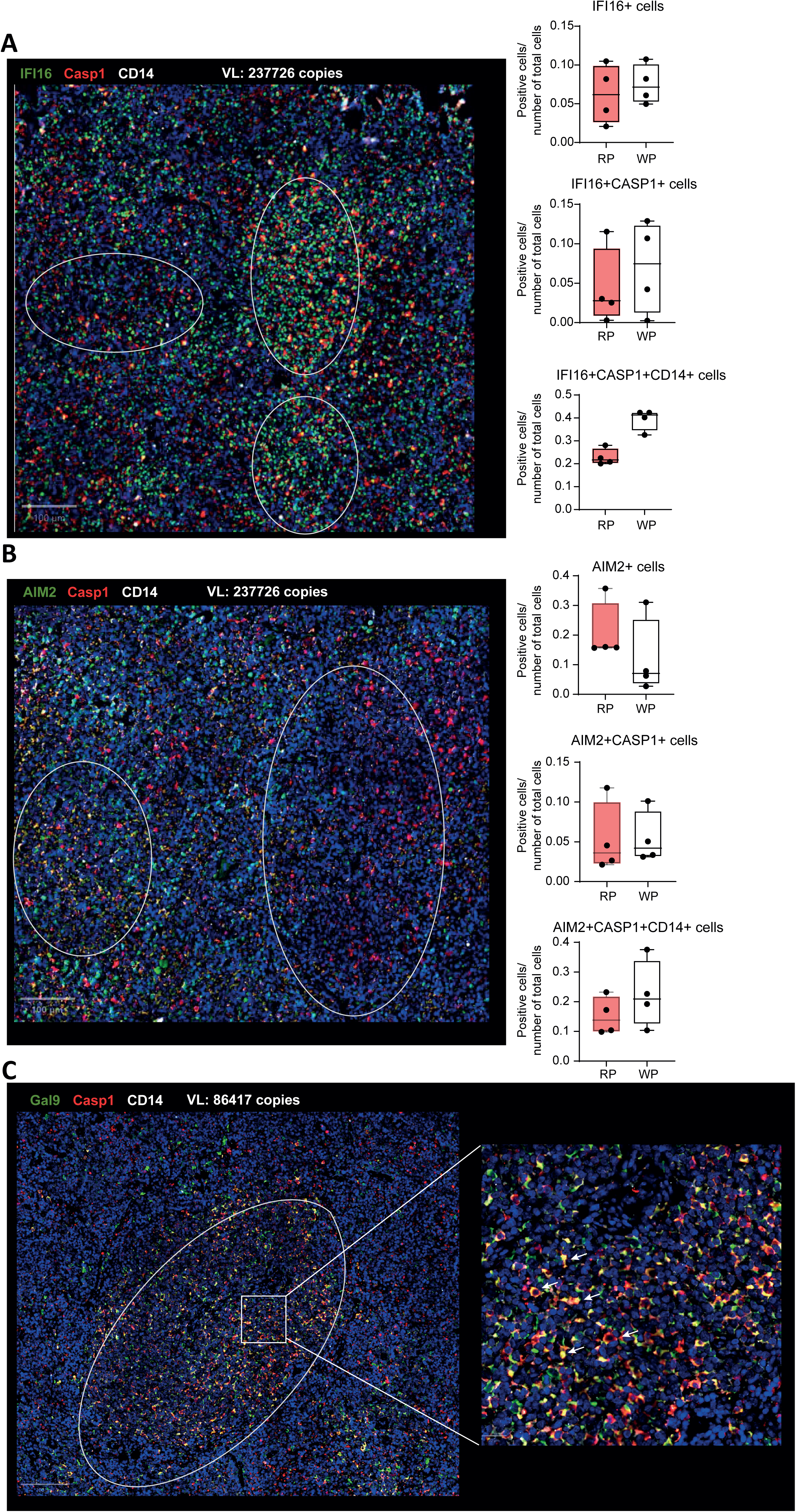
Histological analysis of the inflammasome sensors and ligands expression in myeloid cells from viraemic HIV-1 infected humanized BLT mice. (A-B): Histological analysis of IFI-16 (A) or AIM2 (B) (green) and Caspase-1 (red) expression and DAPI (blue) in the spleen from a viremic HIV-1 infected humanized BLT mouse. Quantifications of cells positive cells for each inflammasome sensor alone (top plots), co-expressing Caspase 1 (CASP1) (middle plots) or both Caspase 1 and CD14 (bottom plots) are shown on the right. (C): Histological analysis of Gal-9 (Gal9; green), CD14 (white) and Caspase-1 (Casp1, red) expression and DAPI (blue) in the spleen from a viremic HIV-1 infected humanized BLT mouse. Zoomed area demonstrating co-expression between Gal9, CD14 and Caspase-1 are shown on the right. Arrows highlight cells co-expressing the mentioned markers

### IFI16/AIM2 inflammasomes control PVR and Gal-9 expression in MDDC from PWH

Since Mo are the source of MDDC used for immunotherapies directed to boost HIV-1 specific CD8+ T cell responses, we asked whether both proinflammatory profiles and higher susceptibility to induce expression of ligands of checkpoint receptors could also be increased in MDDC derived from Mo from PWH after exposure to different PAMPS. To this end, we recruited an additional group of PWH with high and low CD4/CD8 ratios to our study cohort (Supplemental Table 2). No significant differences were observed in baseline proportions of CD86high MDDC expressing PVR and Gal 9 between MDDC from PWH and HD (Supplemental Figure 4A-B). Surprisingly, we did not observe higher basal levels of proinflammatory cytokines IL-1β, IL-6 and TNFα in MDDC from PWH, which in the case of TNFα and IL6 were significantly lower than HD (Supplemental Figure 4B). However, MDDC from PWH increased expression of PVR and Gal-9 in response to Poly I:C, while LPS or CL097 induced lower levels of these molecules compared to baseline (Supplemental Figure 4C-D). Interestingly, while activated MDDC from PWH with high and low CD4/CD8 ratios similarly induced PVR after Poly I:C stimulation (Figure 4A), higher levels of Gal-9 were more efficiently induced by Poly I:C in cells from low CD4/CD8 ratio PWH (Figure 4B). Notably, MDDC from HD did not induce expression of these ligands upon Poly I:C stimulation (Figure 4A-B). Supporting that the induction of checkpoint receptor ligands is linked to a proinflammatory state of MDDC, levels of intracellular TNFα protein were more significantly increased in MDDC from PWH after Poly I:C, CL097 and LPS PAMP stimulation, in contrast to cells from HD. A similar trend was observed with IL-6, although statistical significance was observed only in the presence of LPS (Supplemental Figure 4E). In contrast, no significant further increase was observed in total intracellular IL1-β expression in MDDC from PWH in response to LPS. (Supplemental Figure 4E).

**Figure 4.**
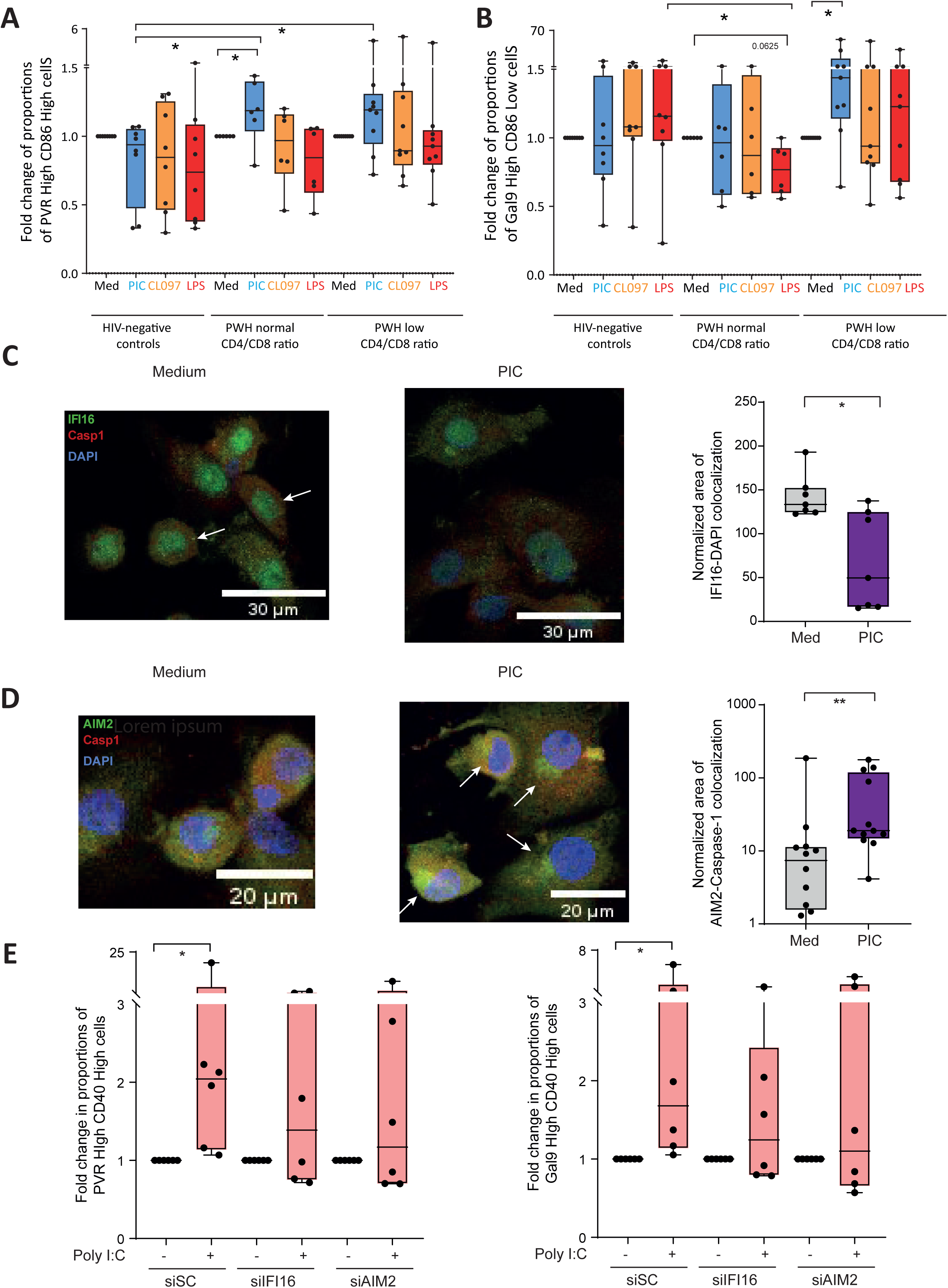
Analysis of dependence on IFI16 and AIM2 inflammasomes regulating induction of PVR and Gal-9 in MDDC from PWH. (A,B): Fold change in proportions of cells expressing high levels of PVR and CD86 (A, left) or high levels of Gal9 with low levels of CD86 (B) in MDDCs cultured for 16h the absence (MED) or presence of Poly I:C (PiC, blue), CLO97, (orange) or LPS (red). (C-D): Confocal microscopy analysis of colocalization of IFI16 (green) with DAPI (blue) (C) or AIM2 (green) with Caspase-1 (D) in MDDC cultured for 16h in the presence of media (grey) or poly IC (violet). Representative images are shown on the left, with the white arrows highlighting the colocalization and quantification of colocalization of IFI-16 with DAPI or AIM2 with caspase-1 is shown on the right. (E): Proportions of MDDC expressing high levels of CD40 and high levels of PVR (left), Gal9 (right) after specific siRNA-mediated knockdown of IFI16 and AIM2 inflammasomes and 16-hour stimulation with Poly I:C. Statistical significance was calculated using two-tailed Wilcoxon test and for the colocalization assays two-tailed Mann-Whitney test was used ( **p<0.01).

To determine whether MDDC from PWH were characterized by higher levels of IFI16 and AIM2 inflammasomes, we analyzed by microscopy the expression of these molecules and caspase 1 in the presence of media or Poly I:C. Similar to Mo, a redistribution of IFI16 from the nucleus to the cytoplasm was observed in MDDC from PWH after Poly I:C stimulation (Figure 4C). However, high levels of colocalization of IFI16 with caspase-1 were already observed in the absence of stimulation suggesting a basal higher predisposition to activate this pathway (Supplemental Figure 4F). Co-distribution of AIM2 and caspase-1 expression in the cytoplasm was significantly higher in MDDC from PWH after Poly I:C stimulation, similarly to our previous observations in Mo (Figure 4D).

To determine whether the observed differences in expression of AIM2 and IFI16 inflammasome sensors could be differentially contributing to the upregulation of ligands for checkpoint receptors in MDDC from PWH after PAMP stimulation, we knocked down the expression of AIM2 and IFI-16 using specific siRNAs (Supplemental Figure 5A). siRNA-mediated knock down of IFI-16 and AIM2 abrogated the upregulation of PVR and Gal-9 in MDDC from PWH treated with Poly IC (Figure 4E) individually or in response to STING agonists or the combination of both stimuli (Supplemental Figure 5B), previously used as adjuvants for MDDC-based immunotherapy^29^. Induction of PVR and Gal-9 expression after stimulation with LPS, flagellin, CL097 was also reduced in Mo after IFI16 siRNA knock down (Supplemental figure 5C). In contrast to MDDC, no significant changes in expression of PD-L1, PVR or Gal-9 were observed in Mo from PWH treated with AIM2-specific siRNAs (Supplemental Figure 5D). Therefore, our data highlight the central role of IFI16 controlling the expression of PVR and Gal-9 in MDDC and Mo from PWH. In contrast, AIM2 contributes to expression of ligands for TIGIT and TIM3 specifically in MDDC from PWH.

### Pharmacological inhibition of the inflammasome potentiates polyfunctional HIV-1 specific CD8+ T cells in vitro

We next asked whether directed inhibition of specific inflammasome components could reduce the induction of ligands of checkpoint receptors in MDDC from PWH treated with Poly IC and STING agonist adjuvant cocktail. Specifically, we tested parthenolide (P; a caspase-1 and NFκB inhibitor), MCC950 (M; NLRP3 inhibitor), a caspase 1-9 inhibitor (V; Z-VAD-FMK) and ODN TTAGGG (O; IFI16/AIM2 inhibitor) in MDDC treated with either individual or combined Poly I:C and STING agonist. We also used BX795 (B; TBK-1 inhibitor) as a control for complete inhibition of MDDC maturation in response to Poly I:C and STING agonists. As expected, Parthenolide and BX795 completely prevented both the induction of mature CD40hi MDDC and the expression of PVR and Gal-9 in these cells (Supplemental Figure 6A); the percentage of CD40Hi cells remained above baseline levels in MDDC treated with Z-VAD-FMK despite a mild significant decrease in these cells in the presence of this drug (Supplemental Figure 6B). As shown in Figure 5A, proportions of activated CD40hi co-expressing PVR and Gal-9 were more efficiently reduced in the presence of Z-VAD-FMK and ODN inhibitors. On the other hand, maturation and expression of checkpoint receptor ligands were not as consistently affected in MDDC cells treated with MCC955 and Parthenolide (Supplemental Figure 6A). Interestingly, no significant differences in intracellular levels of TNFα, IL-6 and IL1β after treatment with Z-VAF-FMK inhibitor (Supplemental Figure 6B). Collectively, these data indicate that the induction of PVR and Gal-9 ligands can be prevented in MDDC from PWH using pharmacological inhibitors of caspases and IFI16/AIM2 sensors.

**Figure 5.**
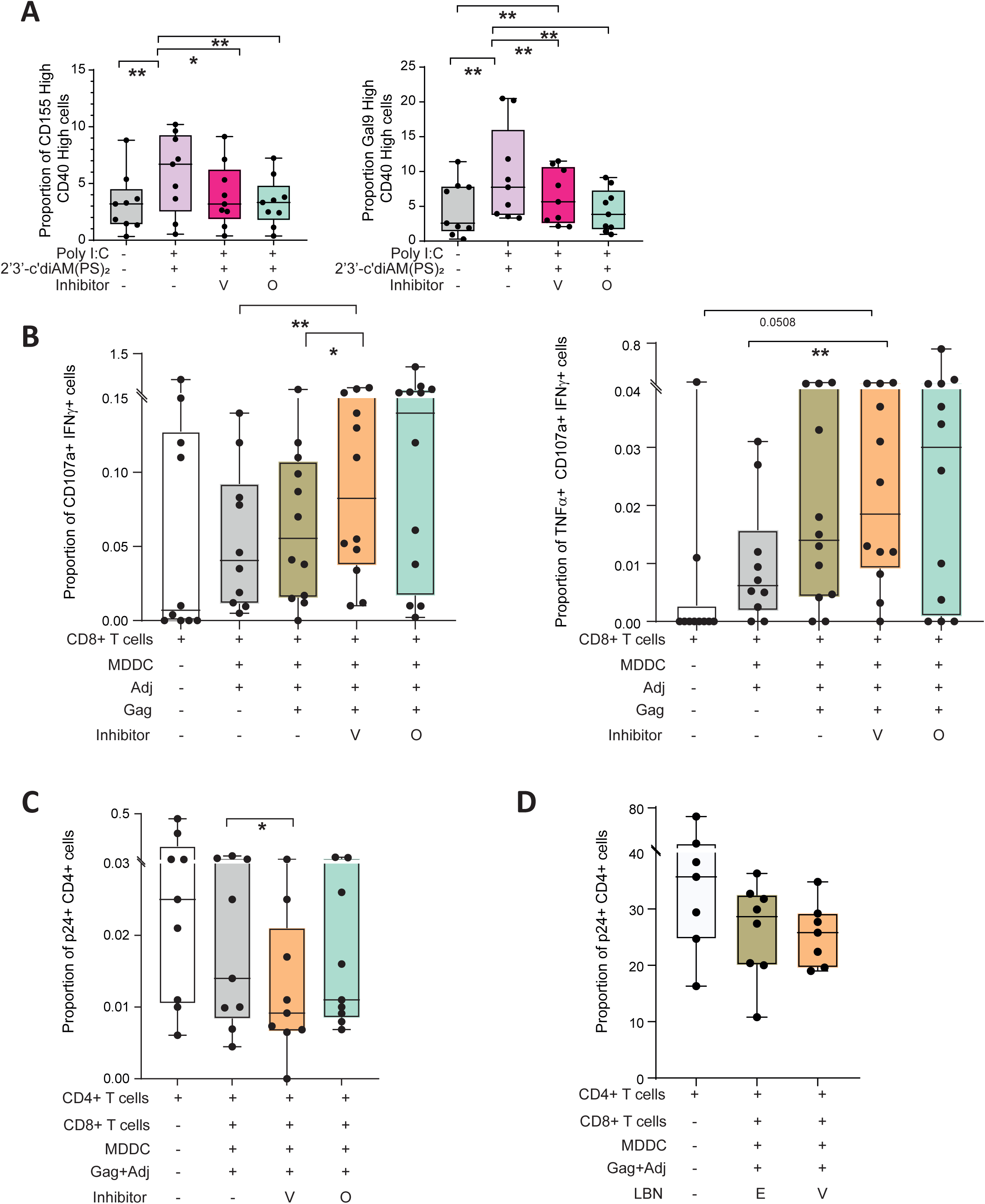
Functional impact of modulation of inflammasome pathways by inhibitors in MDDC on the ligand expression and T lymphocytes-specific responses. (A): Proportion of MDDC expressing high levels of CD40 and high levels of PVR (left) and Gal9 (right) in medium (grey), in presence of Poly I:C and 2’3’-c-diAM(PS)2(Rp,Rp) (STING agonist) alone (pink) or with Z-VAD-FMK (fuchsia) or ODN TTAGGG (mint). (B): Proportions of CD8+ T lymphocytes co-expressing IFNγ and CD107a (left) or in combination with TNFα and (right) alone (white) or after coculture with MDDC treated with Poly I:C and 2’3’-c-diAM(PS)2(Rp,Rp) (STING agonist, Adj) alone (grey), in presence of HIV Gag peptide alone (dark olive) or with Z-VAD-FMK (orange) or ODN TTAGGG (mint). (C): Proportions of p24+ CD4+ T lymphocytes (white) alone, in presence of MDDC activated in presence of Poly I:C and 2’3’-c-diAM(PS)2(Rp,Rp) (STING agonist) and Gag peptide (Gag+Adj) and CD8+ T lymphocytes alone (grey) or in presence of Z-VAD-FMK (orange) or ODN TTAGGG (mint). (D): Proportions of infected p24+ within human C D45+ CD4+ T cells present in spleen of humanized NSG mice transplanted vía i.v. with either only CD4+ T cells from PWH alone (white) or in combination with autologous CD8+ T cells and activated MDDCs and receiving by i.p. injection either empty anti-CD64 mAb-decorated LBN (E, khaki) or Z-VAD-FMK-loaded anti-CD64 mAb-decorated LBN (V, orange). Statistical significance was calculated using two-tailed Wilcoxon test (*p<0.05; **p<0.01).

We then evaluated whether reduction of expression of PVR and Gal-9 ligands with pharmacological inhibition of the inflammasome may improve the efficacy of MDDC-based vaccines stimulating HIV-1 specific CD8+ T cell responses in PWH. To address this possibility, we performed *in vitro* functional assays with MDDC loaded with HIV-1 gag peptides and stimulated with Poly IC and STING agonist adjuvants in the absence (DMSO) or presence of Z-VAD-FMK caspase or ODN TTAGGG IFI16/AIM2 inhibitors in the presence of autologous CD8+ T cells from n=9 PWH with a median expression of 14.1% (min-3.64-27.8) of TIGIT and 1.07% (min 0.06%-max 9.73%) of TIM3. After 16h, we evaluated by FACS the secretion of multiple cytokines such as IFNγ, TNFα and the degranulation marker CD107a by cultured CD8+ T cells (Supplemental Figure 6C). As shown in Figure 5B, proportions of polyfunctional CD107a+ IFNγ+ (left) as well as CD107a+ IFNγ+ TNFα+ (right) HIV-1-specific CD8+ T cells significantly increased in the presence of MDDC treated with Z-VAD-FMK caspase inhibitor. A similar but not significant trend was observed for total IFNγ+ TNFα+ T cells in these assays (Supplemental Figure 6D). Finally, to determine whether increased activation of polyfunctional HIV-1 specific CD8+ T cells translated into more effective elimination of infected cells, we performed *in vitro* co-cultures in the presence of autologous CD4+ T cells treated with the latency reversal agent Romidepsin and in the presence of Raltegravir, to prevent viral expansion. Under these conditions we determined proportion of intracellular p24+ cells included within cultured CD4+ T cells (Supplemental Figure 6E). As shown in Figure 5C, increased reduction of p24+ HIV-1 infected CD4+ T cells was observed in the presence of CD8+ T cells and MDDC treated with Z-VAD-FMK and ODN inhibitors. Finally, we asked whether Z-VAD-FMK inhibitor directed to myeloid cells could enhance immunotherapy against HIV-1. CD64 is highly expressed in Mo and also in MDDC (Supplemental Figure 7A) To this end, we first generated liposome-based nanosystems (LBN) decorated with anti-CD64 mAbs (Supplemental Figure 7B) and tested specific binding to monocytes *in vitro* (Supplemental Figure 7C). Using an *in vitro* analysis of HIV-1 specific response of CD8+ T cell from PWH in the presence of HIV-1 gag peptide stimulation, we confirmed that Z-VAD-FMK similarly induced IFNγ, CD107a expression and tended to boost TNFα secretion on Ag-stimulated CD8+ T cells from PWH (Supplemental Figure 7D). We then tested the efficacy of this treatment boosting the effect of MDDC-CD8+T cell-based immunotherapy *in vivo*. To this end, a total of n=22 NSG mice were injected with CD4+ T cells alone (n=7) or in combination with autologous activated and HIV-1 gag peptide loaded MDDC and CD8 T cells (n=15) of which n=8 received empty and n=7 Z-VAD-FMK loaded anti-CD64 decorated LBN by i.p. injection, which was repeated after 3 days (Supplemental Figure 7E-F). At day 7 of treatment animals were sacrificed and analysis of proportion of p24+ cells was analyzed in human CD4+ T cells present in the spleen from NSG mice (Supplemental Figure 7E-F). As shown in Figure 5D, treatment of animals with MDDC-CD8+ T cells reduced proportions of human HIV-1 infected p24+ cells in the spleen and lower median frequencies of these cells were observed in animals treated with LBN loaded with Z-VAD-FMK. Therefore, inhibition of the + IFI-16/AIM2 inflammasomes improves the ability of MDDC to boost functional HIV-1 specific CD8+ T cell responses and may be a promising modulator of immunotherapy against HIV-1.

## DISCUSSION

DC-based therapies have yielded variable results potentiating immune responses against HIV-1 in PWH, potentially due to an exhausted T cell phenotype linked to chronic HIV-1 infection, which is not fully restored by ART and viral suppression, impairing the capabilities of these cells to respond to immunotherapies ^61–64^. These dysfunctional states have been associated with chronic inflammation and persistent low CD4/CD8 ratio, either by inability to fully restore CD4+ T cell counts or by CD8+ T cell expansion. While most studies have focused on improving immunotherapies by blocking checkpoint receptors expressed by T cells, effectiveness of MDDC may be influenced by inflammatory environment in these PWH individuals. Hence, in this study we hypothesized that low CD4/CD8 T cell ratio would be associated with phenotypically different state of myeloid cells in the context of HIV-1 infection, impairing their abilities to efficiently activate HIV-1 specific CD8+ T cells. In the present study, we have shown that PVR and Gal-9 are differentially induced at the protein level in Mo and MDDC from PWH with low CD4/CD8 T cell ratio, providing novel evidence of a differential inflammatory state in these cells potentially induced by systemic inflammation in this PWH group. Notably, we detected a significant enrichment of PWH with low but detectable viral loads in the PWH group of low CD4/CD8 ratio. Although we observed higher inducibility of PVR and Gal-9 in cells from aviremic PWH with low CD4/CD8 ratios, additional studies should be conducted to completely discard the impact of residual viremia and inflammatory cytokine profiles of these individuals.

Unexpectedly, we have observed higher basal levels of CD40 in Mo from PWH with normal compared to low CD4/CD8 T cell ratios. CD40 is regulated by multiple factors including type II IFN as well as NFkB signaling downstream cytokine receptors and TLR^65,66^ and has been associated to effective IFN-I responses^67^.However, considering lower basal levels of CD40 on these Mo, further studies should determine whether higher activation leads to selective depletion of activated cells from the blood or whether the observed phenotype correspond to a specific trained state in Mo from this group. Therefore, mechanisms involved in CD40 regulation should be investigated in more detail in these PWH groups.

In contrast, plasma levels of IL-1β were higher in PWH with low CD4/CD8 T cell ratio, suggesting a specific and different basal proinflammatory state involving inflammasome activation in this PWH subgroup. We observed that these ligands were induced *ex vivo* by different bacterial and viral PAMPs, but induction was more efficient in activated Mo as well as MDDC from PWH with low CD4/CD8 T cell ratios than in PWH with normal CD4/CD8 T cell ratio. The expression of the main TIGIT Ligand PVR has been described to be regulated by several pathways and mechanisms including NF-kB and TLR through TRIF and MYD88, IL-4 through STAT6, IL-6, IL-22, stress-associated signals and microRNAs^68–73^. Our findings suggest that while PVR is more efficiently induced by Mo from PWH with low CD4/CD8 ratios, MDDC from both PWH groups efficiently induced expression of this molecule. These differences may be related to differential regulatory mechanisms in these two myeloid cell types or by additional activation induced during the differentiation process of MDDC from Mo in the presence of GM-CSF and IL-4. The other checkpoint receptor ligand analyzed in our study is Gal −9, which has been proposed to be induced by type I and II IFNs^74,75^, or by Poly I:C in mesenchymal stem cells^76^. Expression of Gal-9 is also regulated by PI3K/AKT/mTOR pathways^77^. In contrast to PVR, we did consistently observe higher upregulation of this molecule in both Mo and MDDC from PWH with low CD4/CD8 ratios, suggesting conserved mechanisms associated with the inflammatory state of these individuals. Consistent with these observations, transcriptional and protein induction of TNFα, IL-6 was more consistently induced by Mo and MDDC from PWH with low CD4/CD8 T cell ratios in our study, and production of these cytokines may be a biomarker of higher susceptibility to induce TLR signaling, which could contribute to local inflammation in different tissues.

Finally, the ability of Mo and MDDC from PWH to upregulate PVR and Gal-9 as well as their inflammatory profiles after PAMP stimulation associates with increased detection of IL-1β in plasma and by higher expression and activity of IFI16 and AIM2 inflammasomes. Previous studies have already described the association between HIV- 1 sensing by Mo with specific activation of the NLRP3 inflammasome and caspase-1^78,79^. Other cell types, such as CD4+ T cells have been associated with NLRP3 inflammasome induction in HIV-1 infection, inducing Caspase-1 activity after viral antigens recognition or ROS production^58^. In this regard, TLR8 sensing also leads to NLRP3 and induces chromatin remodeling in pro-inflammatory genes such as TNF or IL- 6 in Mo^80^. Although no evident differences were observed in our PWH cohort regarding markers of bacterial translocation such as sCD14 and endotoxin, we observed a correlation with IL-1β levels in plasma consistent with the association between TLR4 signaling with NLRP3 activation in other pathologies^81^. However, in our study we have observed that IFI16 may be an inflammasome whose expression and activity may be higher in Mo and MDDC from PWH, and particularly, in individuals with low CD4/CD8 T cells ratios. Indeed, IFI16 inflammasome has been described to have a role in sensing of different DNA viruses and retroviruses inducing cytoplasmic dsDNA such as, HIV-1 and also controlling CD4+ T cell depletion in the absence of treatment^52–54,82,83^ and its levels were reduced in PWH after ART ^84^. Additional studies using a monocytic cell line (THP-1) described increased expression of IFI16 after HSV-1 infection or after treatment with type-I and type-II IFN^85,86^. In addition, our results suggest that IFI16 may specifically control expression of PVR in both Mo and MDDC after PAMP stimulation. Thus, future studies should determine whether this mechanism could be associated with innate detection of HIV-1 DNA in Mo from PWH. In this regard, a previous study suggested the role of Gal-9 in regulating HIV-1 transcription and viral production in PWH on ART^87^.

Furthermore, we have observed that AIM2 inflammasome may play a more relevant role in MDDC than in Mo regulating PVR and Gal-9. However cellular distribution and colocalization with caspase-1 may be different according to our data. On the one hand, we have shown higher colocalization of IFI16 with caspase-1 in Mo stimulated with Poly I:C. On the other hand, in the case of MDDCs from PWH we observed high colocalization with caspase-1 at baseline and we were able to observe significant changes in the translocation of IFI16 from nucleus to the cytoplasm as a sign of activation^88–90^. In addition, we have observed a significant colocalization of IFI16 and caspase-1 in the nucleus, as suggested by previous studies.^88,91–94^. A possibility is that IFI-16 becomes more activated in MDDC, as a result of differentiation from Mo. In contrast, colocalization of AIM2 with caspase-1 is clearly more efficiently induced by Poly I:C stimulation in MDDC than in Mo from PWH, suggesting that this pathway may differentially contribute to function of specific myeloid cell types. Relevance of these observations was confirmed in tissue from humanized BLT mice, in which we identified different patterns of IFI16 and AIM2 expression upon HIV-1 infection. While IFI16 is mainly localized and co-expressed with caspase-1 in the white pulp of infected mice and is much less presented in red pulp, the expression of AIM2 showed completely inverted pattern. Therefore, these two sensors may be playing different roles in specific myeloid cell types in relevant tissues during infection. Supporting this interpretation, IFI16 is associated both to DNA sensing and to the induction of IFN-β through the activation of TBK1 and IRF3^88,95^. In contrast, AIM2 regulates caspase-1-dependent processing of proinflammatory cytokines in response to DNA but does not mediate type I interferon responses^96^. Nevertheless, AIM2 activation also demonstrated the activation of NF-kB pathway ^97^. Interestingly, IFI16 is involved in the inhibition of caspases activation by AIM2 and NLRP3^98,99^. However, we have not addressed the impact of IFI16 on AIM2 activation in this study. As activation of MDDC and the expression of the costimulatory molecules could affect the efficiency of the antigen presentation^100,101^ we explored the potential of inflammasome inhibitors directed to caspases and IFI16/AIM2 to maintain the expression levels of costimulatory molecules such as CD40 while reducing the levels of PVR and Gal-9. We found that Z-VAD-FMK inhibitor resulted in better efficiency than ODN, more specific IFI16/AIM2 sensing. These results could be explained by a broader effect of other inflammasome pathways than just IFI16/AIM2 inflammasome regulating PVR and Gal-9, or that the IFI16/AIM2 inhibitors also interfere with physiological activation of MDDC, therefore additional *in vitro* studies should be performed.

Nevertheless, we addressed the functional impact of inhibiting IFI16/AIM2 inflammasomes on the ability of MDDCs to activate HIV-specific CD8+ T cell cytotoxicity that resulted in reduction of p24+ CD4+ T cells. Our data show that pre-treatment of MDDC with these inhibitors allow to improve their ability to activate polyfunctional HIV-1 specific CD8+ T cells. These observations are consistent with the idea that inhibition of inflammasomes might improve the immune response against tumors^102,103^, and we have now expanded the potential targets that may be useful for therapeutic modulation. However, further preclinical *in vivo* validation should be performed in humanized animal models.

Collectively, our work highlights the association between systemic inflammation and higher induction expression of Gal-9 and PVR ligands controlled by IFI16/AIM2 inflammasomes in Mo and MDDC from PWH with low CD4/CD8 ratios. Targeted modulation of these sensors could be an attractive strategy to further improve MDDC-based immunotherapy against HIV-1 and boost efficacy of cure strategies.

## AUTHOR Contributions

EMG, MCM and IT designed the study and prepared the manuscript MCM and IT executed most of the experiments and analyses

JPP performed analysis of the induction of checkpoint ligands, cytokine secretion and activation molecules in MDDC of PWH upon stimulation with PAMPs and the impact of Z-VAD-FMK on cytokine secretion

ISC participated in the *in vivo* experiments, microscopy image analysis and provided tissue from HIV-1 infected humanized mice

MAL participated in processing of PB from PWH and also tissue samples and cell isolation for *in vivo* experiments

PNHA participated in PWH sample processing, cell isolation and microscopy experiments

RG participated in the *in vivo* experiments and with OP provided technical support in the study LGF, IS, SSV and SM provided blood samples of PWH recruited at Hospital de la Princesa and Hospital Ramón y Cajal

PLC, ML and JAL manufactured and provided antibody-covered LBN containing fluorochromes or inflammasome inhibitors

FSM and AA provided buffy coat samples and for immune exhaustion reagents and critical feedback during the study

EMG supervised the study

## ACKNOWLEDGMENTS

E.M.G was supported by the Spanish Agencia Estatal de Investigación RETOS, Generación de conocimiento and consolidation programs (PID2021-127899OB-I00; CNS2023-144841; PID2024-160973OB-I00), GLD24/00117 grant from Gilead biosciences, La Caixa Banking Foundation ETI-CureHIV (HR20-00218), MORELIA (ICI20/00058) and infectious diseases CIBER from ISCIII (CB21/13/00107). M.C.M was supported by La Caixa Banking Foundation ETI-CureHIV (HR20-00218). I.T. was supported by FPI UAM fellowship. R.G. was supported by PID2021-127899OB-I00 and GLD24/00117 grants. M.A.L was supported by the Formación de Personal Investigador (FPI) grant PRE2022-104516. P2022/BMD7209-INTEGRAMUNE from Comunidad Autónoma de Madrid and La Caixa Health Research Grant LCF/PR/HR23/52430018 and PID2023-149541OB-I00 to F.S.M also supported the study. I.S.C was supported by infectious diseases CIBER from ISCIII (CB21/13/00107). Optical microscopy was conducted at the Video-microscopy Facility of the IIS-Princesa (Madrid, Spain) co-funded by IFEQ21/00085 and IFCS22/00014 from Instituto de Salud Carlos III (ISCIII) and FEDER. SSV reports funding by Instituto de Salud Carlos III (PI21/00041, and PI24/00078 to SSV) and the European Union (NextGenerationEU); CIBERINFEC IM23/INFEC/4 to SSV) and CIBER (iN24-01). P.N.H.A.was supported by MORELIA (ICI20/00058) grant. PLC reports funding by the Consejería de Economía, Conocimiento, Empresas y Universidad de la Junta de Andalucía-Universidad de Sevilla (VII PPIT-FEDER Andalucía–A-2024-31730, Feder Funds and FQM-206) and the Ayuda Servicios Generales de la Universidad de Sevilla, CITIUS, VII PP2025/00000399).

## DICLOSURE AND COMPETING INTERESTS STATEMENT

The authors declare that they have no competing interests.

## SUPPLEMENTAL FIGURES

**Supplemental Figure 1. Plasma markers of inflammation and bacterial translocation in PWH.** (A) Heatmaps reflecting Spearman correlation networks between CD4/CD8 ratio (ratio), years under ART (years), age, CD4 T cell counts at diagnosis (NADIR) and at sample collection (CD4 count) and plasma viral load (VL sample) in all PWH recruited (left) or within low (middle) or normal (right) CD4/CD8 ratio PWH subgroups. (B): Analysis of plasma viral load (mRNA copies/mL) as well as concentrations of soluble CD14 (ssCD14), endotoxin and IL-6 concentration from n=20 PWH with low (n=12) or normal (n=8) CD4/CD8 ratios. (C) Flow cytometry gating strategy for analysis of ligands for checkpoint inhibitory receptors in live large HLADR+ Mo selected by CD14 immunomagnetic sorting from the blood of PWH. (D-E) Proportion of CD40 High Mo (D), CD40 High Gal9+ Mo from aviremic donors (E), CD40 High PVR High Mo from aviremic donors (F) or CD40 High PD-L1 High Mo from all cohort (top) or from aviremic donors only (bottom) (G) in groups of HIV- 1-negative controls, and PWH with low or normal CD4/CD8 T cell ratio after 16-hour stimulation with PAMPs mimicking bacterial translocation (LPS, red; Flagellin, Flag, purple) or viral replication (CL097, ssRNA, orange; Poly I:C, dsRNA, blue). Statistical significance was calculated using Kruskal-Wallis test for comparison of stimulation with the same PAMP between groups (*p<0.05; **p<0.01), and One-Way ANOVA Friedman test with Dunn’s multiple comparison test for comparison within the same group (*p<0.05; **p<0.01; ***p<0.001; ****p<0.0001).

**Supplemental Figure 2. Analysis of expression of cytokines and inflammasome sensors in Mo from PWH with low and normal CD4/CD8 T cell ratios.** (A-B): RT-qPCR analysis of mRNA expression of TNFα, IL6 and IFNβ (A) or NLRP3 and NLRC4 inflammasomes (B) normalized to β-actin in Mo from HIV-1 negative controls, and PWH with low or normal CD4/CD8 T cell ratio after 16-hour stimulation with PAMPs mimicking bacterial translocation (LPS, red; Flagellin, Flag, purple) or viral replication (CL097, ssRNA, orange; Poly I:C, dsRNA, blue). Statistical significance was calculated using Kruskal-Wallis test for comparison of stimulation with the same PAMP between groups (*p<0.05; **p<0.01; ***p<0.001; ****p<0.0001), and One-Way ANOVA Friedman test with Dunn’s multiple comparison test for comparison within the same group (*p<0.05).

**Supplemental Figure 3. Histological analysis of the inflammasome sensors and ligands expression in myeloid cells from HIV-1 infected humanized BLT mice with low or absent viral load.** (A): Histological immunofluorescence analysis of HIV-1 p24 (white) and DAPI (blue) in the spleen from HIV-1 infected hBLT mouse (B-E): Histological immunofluorescence analysis of IFI-16 (B, C), AIM2 (D) or PVR (E) (green), Caspase-1 (red), CD14 (white) expression and DAPI (blue) in the spleen from representative HIV-1 infected hBLT mouse displaying low viral load (B) or uninfected (C) animals. Different zoomed areas corresponding to broad or localized inflammasome sensor expression are shown on the right. Quantification of total CASP1+ and CD14+ CASP1+ cells in red pulp (red, RP) and white pulp (WP, white) in spleen from infected hBLT mice is shown in the lower panel B. Zoomed areas highlighting co-expression between IFI16 (B, C), AIM2 (D), PVR (E) and Caspase-1 are shown on the right.

**Supplemental Figure 4. Analysis induction of PVR and Gal-9 in MDDC from different groups of PWH.** (A): Representative staining and gating strategy determining the expression of PVR, Gal9, TNFα, IL-6 and IL-1β in MDDCs from PWH (B): Basal levels of intracellular expression of TNFα, IL-6 and proportions of cells expressing high levels of CD86 and Gal-9 and PVR in MDDC. (C,D): Fold change in proportions of cells co-expressing high levels of PVR and CD86 (C), and proportions of Gal9hi CD86low (D, left), and Gal9hi CD86hi (D, right) in MDDC from all PWH donors after Poly I:C (PiC, blue), CLO97 (orange) or LPS (red) stimulation. Data was normalized to values present in cultures performed in media (Med). (E) Fold change in proportions of MDDC from HD (upper line) or PWH (lower line) expressing TNFα (left), IL-6 (center), IL-1β (right) alone (-) or upon stimulation with Poly I:C (PiC, blue), CLO97 (orange) or LPS (red). (F): Quantification of the colocalization between IFI16 and Caspase-1 from MDDC in PWH. Statistical significance was calculated using two-tailed Wilcoxon test. (*p<0.05; **p<0.01).

**Supplemental Figure 5. Impact of modulation by siRNA-mediated knockout of inflammasome pathways in MDDC on the ligand expression.** (A): Representative efficiency of siRNA-mediated knockout of IFI16 (n=6) and AIM2 (n=6) evaluated by RT-qPCR analysis compared to control sc-siRNA treated cells. (B): Proportions of MDDC expressing high levels of CD40 and high levels of PVR (left), or Gal-9 (right) after specific siRNA-mediated knockdown of IFI16 and AIM2 inflammasomes and 16-hour stimulation with 2’3’-c-diAM(PS)2(Rp,Rp) (STING agonist) alone or in combination with Poly I:C. (C-D): Analysis of proportions of CD40Hi PVR High (upper plots) and Gal9+ CD40 High (bottom plots) from Mo from PWH nucleofected with either scramble (SC), IFI16-specific (C) or AIM2 (D) siRNAs and cultured for 16h in media or in the presence of LPS (red), Flagellin (violet), CL097 (orange) or Poly I:C (PIC, blue). Statistical significance was calculated using two-tailed Wilcoxon test. (*p<0.05; **p<0.01)

**Supplemental Figure 6. Functional impact of the inhibitors on the modulation of inflammasome pathways in MDDC on the ligand expression and HIV-specific T lymphocyte responses.** (A): Proportion of MDDC expressing high levels of CD40 and high levels of PVR (left) and Gal-9 (right) alone (grey), in presence of Poly I:C and 2’3’-c-diAM(PS)2(Rp,Rp) (STING agonist) alone (pink) or in presence of Parthenolide (orange), MCC950 (turquoise) or BX795 (coral red). (B): Proportions of MDDC from PWH expressing TNFα (left), IL-6 (center) and IL-1β (right) absence of stimuli (grey) or presence of Poly I:C and 2’3’-c-diAM(PS)2(Rp,Rp) (STING agonist) alone (pink) or in presence of Z-VAD-FMK (fuchsia). (C): Flow cytometry gating strategy for analysis of CD8+ T lymphocytes from PWH and the expression of IFNγ, TNFα and CD107a performed in the presence of of MDDCs with HIV-1 Gag peptide stimulation. FMO and PMA+Ionomycine conditions were included as a negative and positive staining control respectively. (D): Proportions of CD8+ T lymphocytes (right) co-expressing TNFα and IFNγ alone (white) or after coculture with MDDC treated with Poly I:C and 2’3’-c-diAM(PS)2(Rp,Rp) (STING agonist) alone (grey), in presence of HIV Gag peptide (dark olive) or with Z-VAD-FMK (blue) or ODN TTAGGG (orange). (E): Flow cytometry gating strategy for analysis of CD4+ T lymphocytes from PWH expressing intracellular HIV-1 p24 in the in vitro functional assays of elimination of infected cells. FMO condition for p24 was included as a negative staining control. Statistical significance was calculated using two-tailed Wilcoxon test. (*p<0.05; **p<0.01).

**Supplemental Figure 7. I*n vivo* assays executed with LBN.** (A): Representative flow cytometry staining of CD64 in human MDDC. (B): Schematic representation of the LBN structure and integration of anti-CD64 mAb decoration (left) and representative transmission electron microscopy image of anti-CD64-covered LBN generated for the study. (C): Representative flow cytometry dot plots showing incorporation of fluorescent control naked CFCL-containing LBN (left plot) or anti-CD64mAb-decorated LBN (right plot) in Mo from PWH. Analysis of proportions of fluorescent Mo incubated with LNP not decorated and without fluorescent probe (white) or containing CFCL alone without mAb decoration (light green) or decorated with different 2.5 μg/ml (acid green) or 10 μg/ml (olive green) of anti-CD64 mAb-decorated LBN are shown in the right (D): Analyses of fold change in proportions of CD8+ T cells expressing CD107a (left), IFNγ,(middle) and TNFα (right) from PWH cultured with autologous activated MDDCs treated with DMSO (white), soluble Z-VAD-FMK (khaki), empty (E, white) or Z-VAD-FMK loaded (orange) anti-CD64 mAb-decorated LBNs. (C): Schematic representation of the *in vivo* assay to test LBN modulation of inflammasome combined with MDDC-CD8+T cell immunotherapy using the humanized mVOA model. (D): Representative FACS gating strategy used to identify live human CD45+ CD3+ CD8- T cells expressing HIV-1 p24 in the spleen of a representative humanized NSG mouse from the *in vivo* assays.

