## Supplementary figures and images for "IFI16/AIM2 inflammasomes control Gal-9 and PVR in myeloid cells from PWH and their targeting improves immunotherapy against HIV-1"

### Supplemental Figure 1

Supplemental figure 1

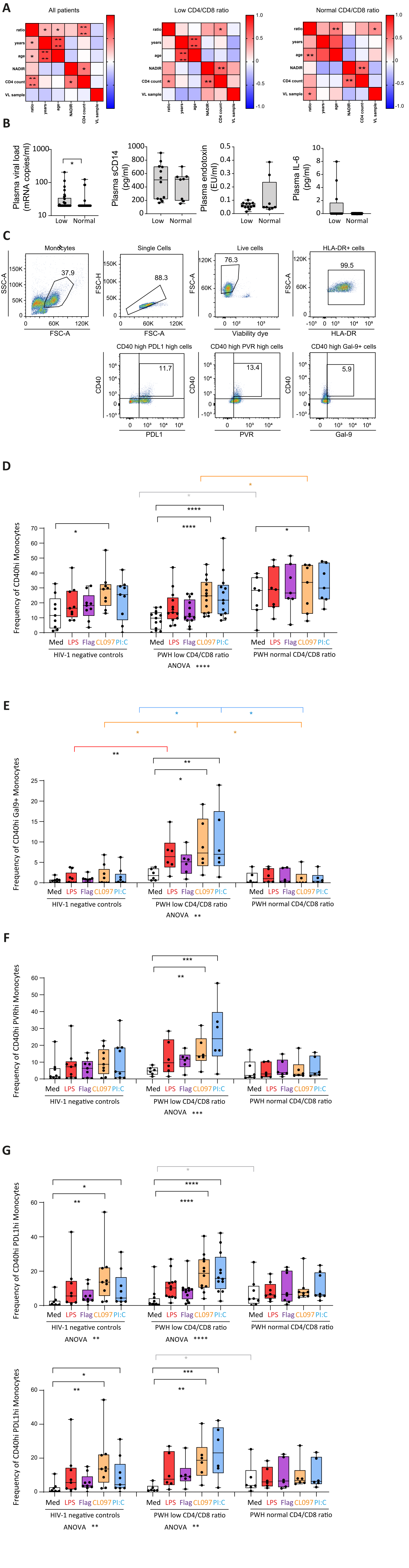

### Supplemental Figure 2

#### Supplemental figure 2

**A**

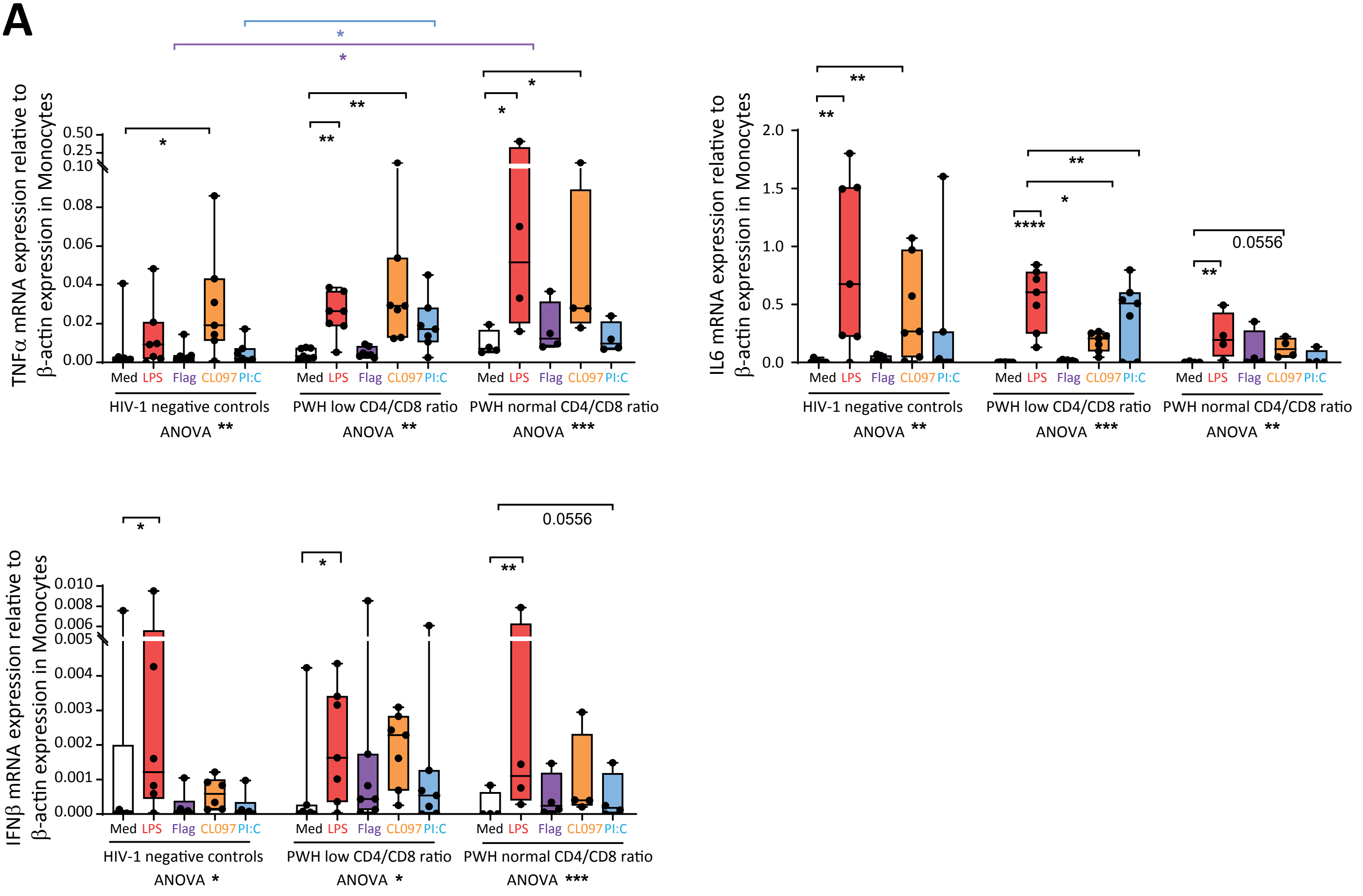


**B**

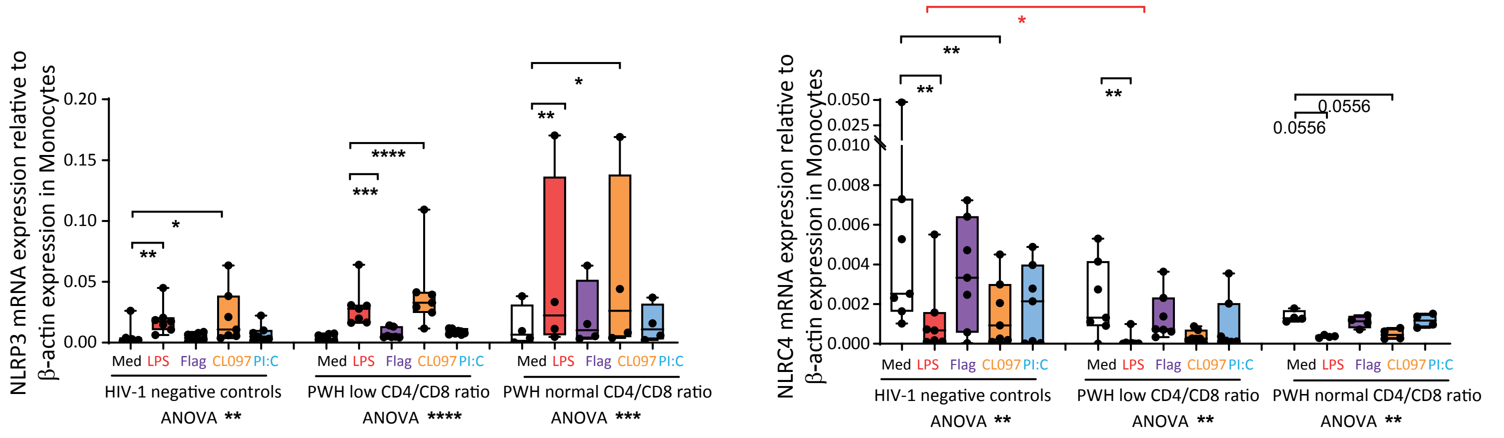

### Supplemental Figure 3

Supplemental figure 3

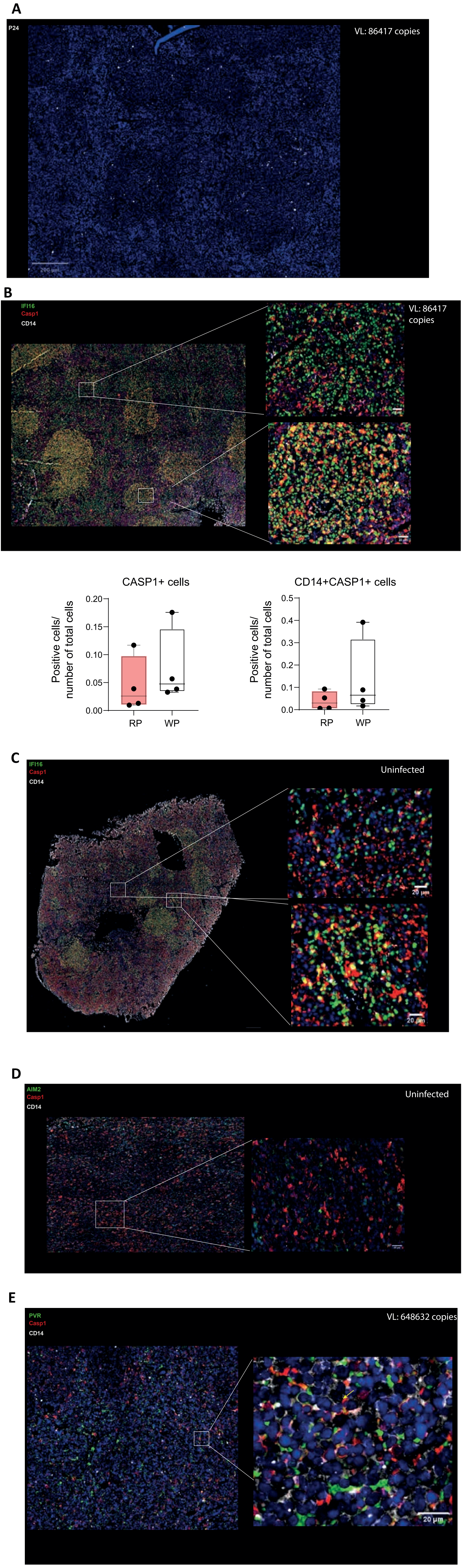

### Supplemental Figure 4

Supplemental figure 4

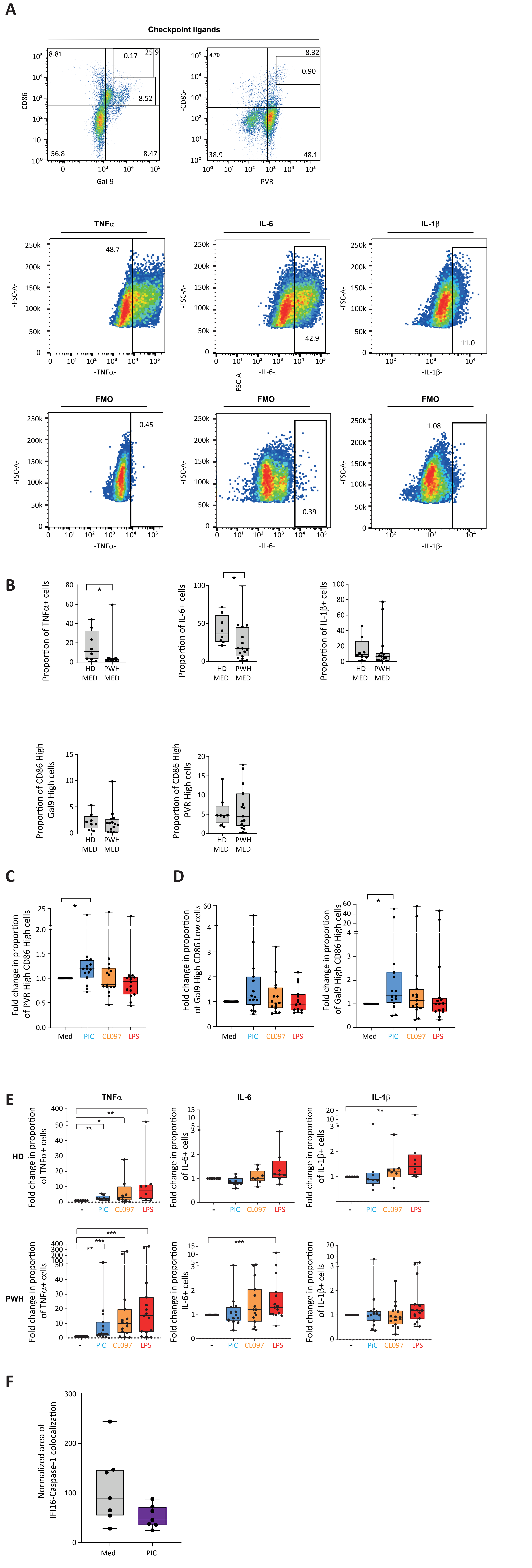

### Supplemental Figure 5

#### Supplemental figure 5

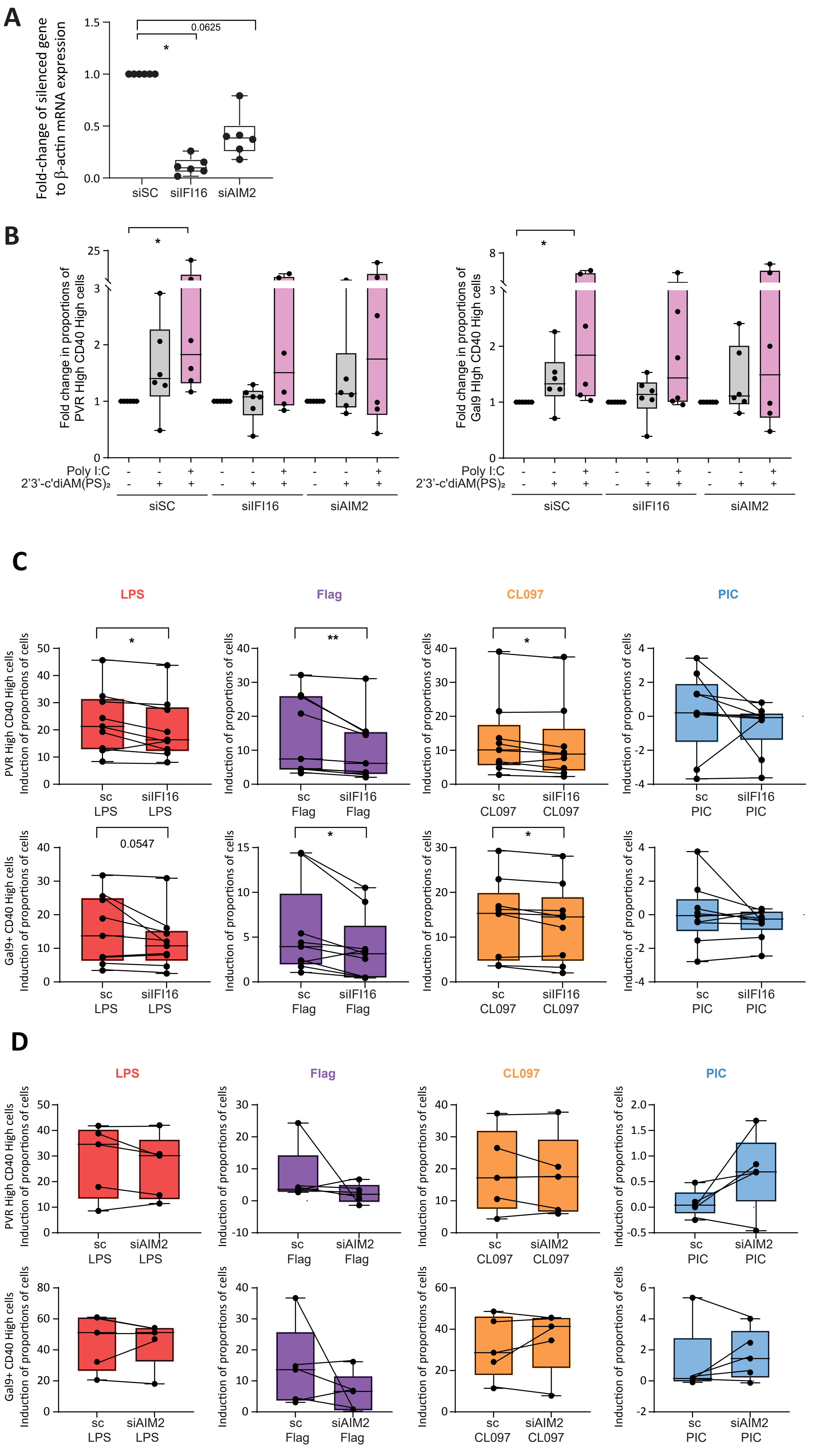

### Supplemental Figure 6

#### Supplemental figure 6

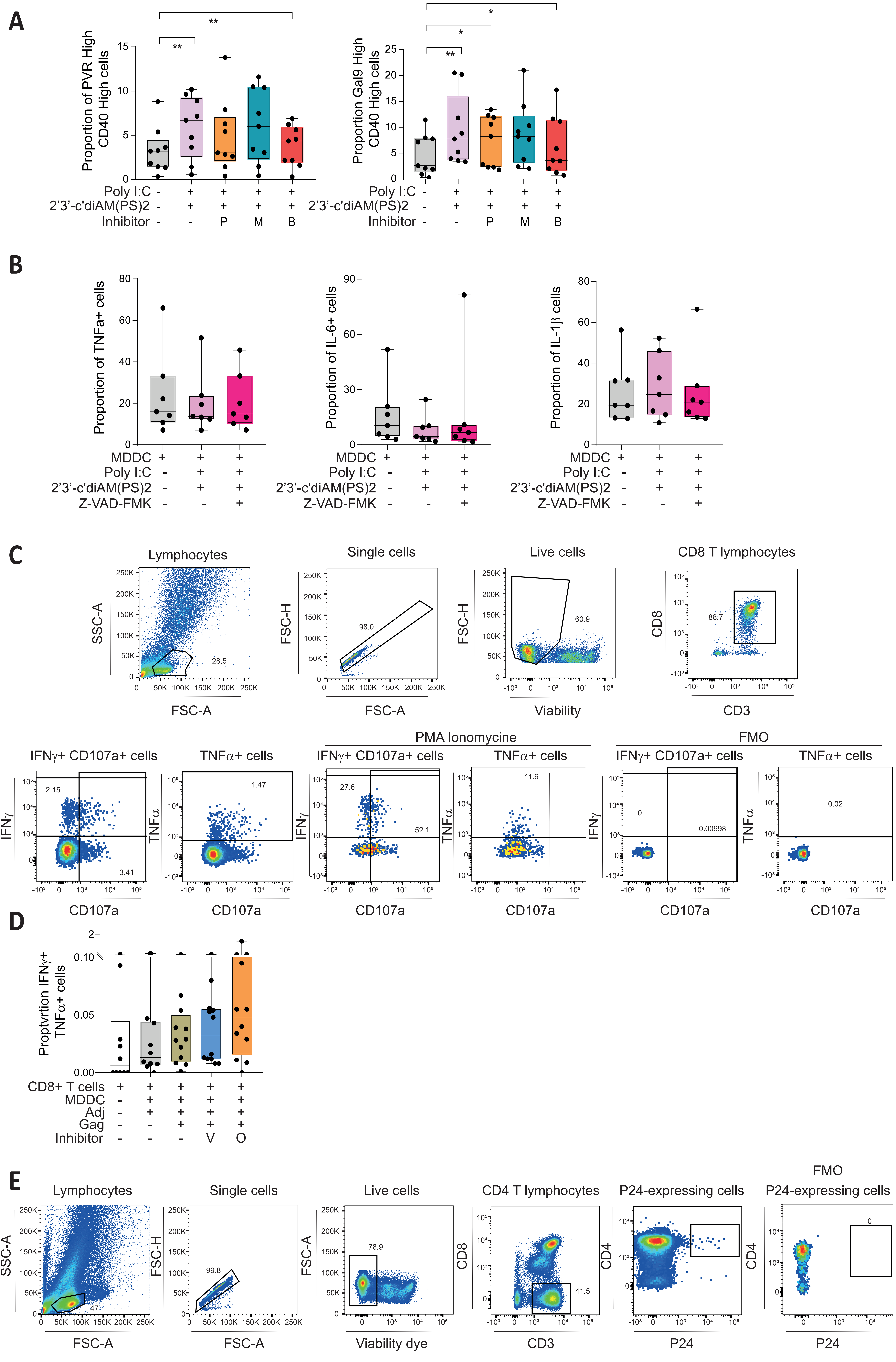

### Supplemental Figure 7

#### Supplemental figure 7

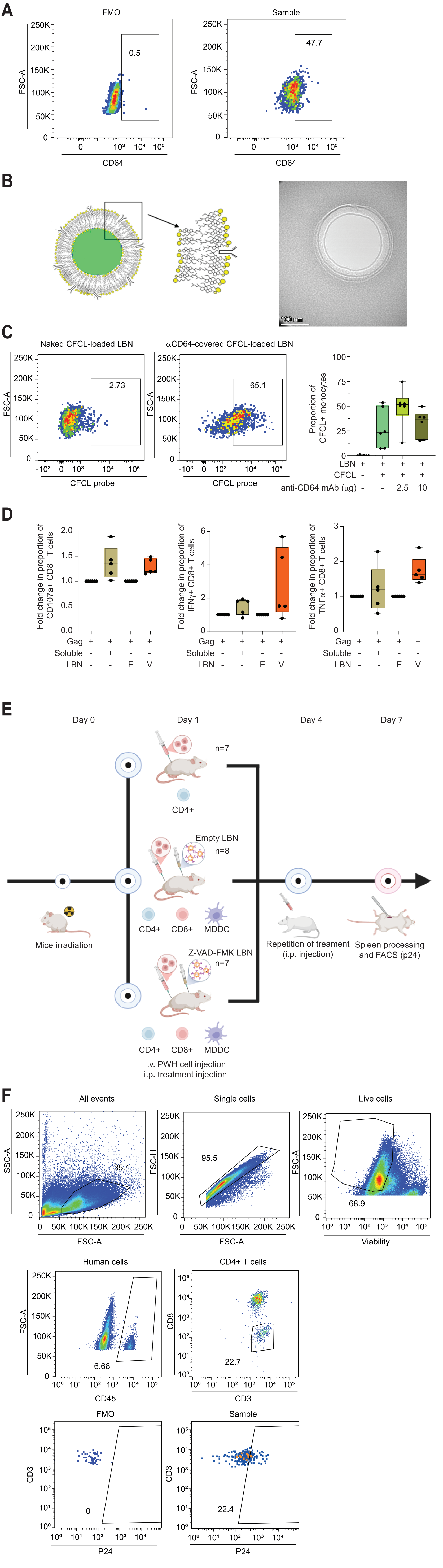
