## Supplemental Tables for "IFI16/AIM2 inflammasomes control Gal-9 and PVR in myeloid cells from PWH and their targeting improves immunotherapy against HIV-1"

|  | Total cohort | Low CD4/CD8 ratio | Normal CD4/CD8 ratio | p-value |
| --- | --- | --- | --- | --- |
| Number of donors, median (min-max) | 60 | 29 | 31 |  |
| Ratio CD4/CD8, median (min-max) | 1.015 (0.16-3.73) | 0.6 (0.16-0.79) | 1.36 (1.01-3.73) | <0,0001 |
| Years on ART, median (min-max) | 9 (1-39) | 7 (1-26) | 10 (1-39) | 0,0771 |
| Age, median (min-max) | 47 (27-73) | 44 (27-71) | 48 (27-73) | 0,3111 |
| Sex, male (%) | 50 (83) | 25 (86) | 25 (81) | 0.5634 |
| NADIR, median (min-max) | 396.5 (1.99-1628) | 370 (1.99-752) | 407 (11-1628) | 0,4987 |
| CD4 count, , median (min-max) | 842 (143.09-1628) | 759 (143.09-1529.67) | 941(338-1628) | 0,0009 |
| Viral load, median (min-max) | 20 (20-205) | 20 (20-205) | 20 (20-124) | 0,0251 |

Supplemental Table 1. Clinical parameters of cohort of PWH involved in study

|  | Total cohort |
| --- | --- |
| Number of donors, median (min-max) | 13 |
| Ratio CD4/CD8, median (min-max) | 0.9 (0.81-3.73) |
| Years on ART, median (min-max) | 9 (2-27) |
| Age, median (min-max) | 45(28-72) |
| Sex, male (%) | 12 (92) |
| NADIR, median (min-max) | 357 (252-531) |
| CD4 count, , median (min-max) | 806.46 (468.72-1365) |
| Viral load, median (min-max) | 20 (20-32) |

Supplemental Table 2. Clinical parameters of the additional cohort of PWH involved in study

| Gene | Sequence |
| --- | --- |
| β-actin | Forward: CTGGAACGGTGAAGGTGACA  Reverse: CGGCCACATTGTGAACTT |
| AIM2 | Forward: CAGACCCGGTTTGCTGATCG  Reverse: CCAGGCCTGTTAGCAAGAGTA |
| IFI16 | Forward: ACTGAAGGAGCAGAGGCAAC  Reverse: TCACTGGGCGTTTTTGGAGA |
| IFNβ | Forward: CTTGGATTCCTACAAAGAAGCAGC  Reverse: TCCTCCTTCTGGAACTGCTGCA |
| IL-6 | Forward: CCTGAACCTTCCAAAGATGGC  Reverse: TTCACCAGGCAAGTCTCCTCA |
| IL-1β | Forward: ATGATGGCTTATTACAGTGGCAA  Reverse: GTCGGAGATTCGTAGCTGGA |
| NLRC4 | Forward: GGCAATTGGATTGCTCAGCC  Reverse: GGAAAGGTCAAAGGTGATCCCA |
| NLRP3 | Forward: GATCTTCGCTGCGATCAACA  Reverse: GGGATTCGAAACACGTGCATTA |
| TNFα | Forward: CAGCCTCTTCTCCTTCCTGAT  Reverse: GCCAGAGGGCTGATTAGAGA |

Supplemental Table 3. Primers used for qPCR analysis

| Antibody | Clone | Catalogue reference | Provider |
| --- | --- | --- | --- |
| Ghost Dye Red 780 |  | 13-0865-T100 | Tombo biosciences |
| LIVE/DEAD Fixable Yellow 405 |  | L34959 | Invitrogen |
| CD3 CF-Blue | 33-2A3 | 3CFB1-100T | Immunostep |
| CD3 PE/Cyanine7 | HIT3a | 300316 | Biolegend |
| CD3 FITC | HIT3a | 300306 | Biolegend |
| CD4 APC | A161A1 | 357408 | Biolegend |
| CD4 Pacific Blue | OKT4 | 317429 | Biolegend |
| CD8a PerCP | HIT8a | 300922 | Biolegend |
| CD8 PeCY7 | RPA-T8 | 557746 | BD |
| CD11c Pacific Blue | 3.9 | 301626 | Biolegend |
| CD14 PercP | 63D3 | 367110 | Biolegend |
| CD40 FITC | 5C3 | 334306 | Biolegend |
| CD86 PE/Cyanine7 | BU63 | 374210 | Biolegend |
| CD86 APC R700 | 2331 (FUN-1) (RUO) | 565149 | BD |
| CD107a APC | H4A3 | 328620 | Biolegend |
| CD155 PE/Cyanine7 | SKII.4 | 337614 | Biolegend |
| Galectin-9 PerCP/Cyanine5.5 | 9M1-3 | 348910 | Biolegend |
| HIV-1 p24 PE | KC57-RD1 | 6604667 | Beckman coulter |
| HLA-DR Pacific Blue | L243 | 307633 | Biolegend |
| IFNγ FITC | 4S.B3 | 554551 | BD |
| PD-L1 Brilliant Violet 711 | 29E.2A3 | 329722 | Biolegend |
| TNFα Pacific Blue | Mab11 | 502920 | Biolegend |
| TNFα PercP | Mab11 | 502924 | Biolegend |
| IL-6 APC | MQ2-13A5 | 501112 | Biolegend |
| IL-1β FITC | H1b-98 | 511705 | Biolegend |

Supplemental Table 4. Antibodies used for FACS analysis

| Antibody | Host species | Clone | Reference | Provider |
| --- | --- | --- | --- | --- |
| IFI16 | Rabbit | Polyclonal | Ab169788 | abcam |
| Caspase-1 | Goat | Polyclonal | AF6215 | R&D |
| AIM2 | Rabbit | Polyclonal | Ab233033 | abcam |
| CD14 | Mouse | 4B4F12 | Ab182032 | abcam |
| PVR | Rabbit | Polyclonal | 81254S | Cell signaling |
| Anti-rabbit AF488 | Donkey | Polyclonal | R37118 | Invitrogen |
| Anti-rabbit AF488 | Donkey | Polyclonal | A21206 | Invitrogen |
| Anti-mouse AF647 | Donkey | Polyclonal | A31571 | Invitrogen |
| Anti-goat AF568 | Donkey | Polyclonal | A11057 | Invitrogen |

Supplemental Table 5. Antibodies used for histological analysis
